# An RNA-binding switch drives ribosome biogenesis and tumorigenesis downstream of RAS oncogene

**DOI:** 10.1101/2021.12.16.472890

**Authors:** Muhammad S. Azman, Martin Dodel, Federica Capraro, Rupert Faraway, Maria Dermit, Wanling Fan, Jernej Ule, Faraz K. Mardakheh

## Abstract

Oncogenic RAS signaling reprograms gene expression through both transcriptional and post-transcriptional mechanisms. While transcriptional regulation downstream of RAS is relatively well characterized, how RAS post-transcriptionally modulates gene expression to promote malignancy is largely unclear. Using quantitative RNA Interactome Capture analysis, we reveal that oncogenic RAS signaling reshapes the RNA-bound proteomic landscape of cancer cells, with a network of nuclear proteins centered around Nucleolin displaying enhanced RNA-binding activity. We show that Nucleolin is phosphorylated downstream of RAS, which increases its binding to pre-ribosomal-RNA (rRNA), boosts rRNA production, and promotes ribosome biogenesis. This Nucleolin-dependent enhancement of ribosome biogenesis is crucial for RAS-induced cancer cell proliferation, and can be targeted therapeutically to inhibit tumor growth. Our results reveal that oncogenic RAS signaling drives ribosome biogenesis by regulating the RNA-binding activity of Nucleolin, and highlight the crucial role of this process in RAS-mediated tumorigenesis.

## Introduction

*RAS* genes are amongst the most mutated proto-oncogenes in human cancers, with around 20% of all malignancies estimated to harbor oncogenic mutation in one of the three highly homologous *KRAS, NRAS*, or *HRAS* genes (Prior *et al*, 2020). The encoded RAS proteins are small GTPases which function as molecular switches that regulate several downstream kinase signaling pathways, including ERK1/2 (also known as the RAS-MAPK cascade), and PI3K (Malumbres & Barbacid, 2003). Oncogenic mutations in RAS result in constitutive activation of these downstream kinase signaling pathways, which phosphorylate a plethora of cellular substrates, including a number of transcription factors, ultimately resulting in significant changes in the gene expression profile that promote various aspects of malignancy (Pylayeva-Gupta *et al*, 2011). Although this transcriptional regulation has been shown to play a key role in RAS tumorigenesis, recent studies have revealed that a significant degree of gene expression dysregulation that occurs in cancers is post-transcriptional (Nusinow *et al*, 2020). However, how RAS signaling post-transcriptionally modulates gene expression remains largely unclear.

RNA-binding Proteins (RBPs) are the main post-transcriptional regulators of gene expression, controlling all aspects of the RNA life-cycle from synthesis to degradation (Gerstberger *et al*, 2014). Numerous studies have revealed a key role for many RBPs in cancer development and progression (Kang *et al*, 2020). These RBPs coordinate diverse aspects of post-transcriptional regulation, including splicing (Fish *et al*, 2021), post-transcriptional modifications (Barbieri *et al*, 2017), transport (Dermit *et al*, 2020), translation (Truitt *et al*, 2015), and turnover of various types of RNA (Yu *et al*, 2020). The recent advent of RNA Interactome Capture (RIC) methods, which allow global unbiased identification of proteins that are directly bound by RNA *in vivo,* has transformed our understanding of RBPs (Baltz *et al*, 2012; Castello *et al*, 2012; Queiroz *et al*, 2019; Trendel *et al*, 2019). RIC studies have significantly expanded the catalogue of known RBPs, with around 10% of the human proteome having been demonstrated to bind RNA. These include not only ‘conventional’ RBPs, which contain at least one classical globular RNA-binding Domain (RBD), but also ‘non-conventional’ RBPs that lack any apparent RBDs (Hentze *et al*, 2018). When combined with quantitative proteomics, RIC can also be used for quantifying changes in the RNA-bound proteome (RBPome), thus allowing systematic study of RNA-binding dynamics (Garcia-Moreno *et al*, 2019; Sysoev *et al*, 2016). However, a comprehensive understanding of how oncogenic signaling pathways dynamically modulate the RBPome is still lacking.

Here, we devised a quantitative whole-transcriptome RIC approach, in order to define the impact of oncogenic RAS signaling on the RBPome of mouse Pancreatic Ductal Adenocarcinoma (PDAC) cells. We focused on PDAC, as nearly all its cases harbor oncogenic *KRAS* mutations, highlighting a key role for RAS in the etiology of the disease (Waters & Der, 2018). Our results reveal that through activation of downstream Erk1/2 signaling, oncogenic Kras extensively remodels the RBPome of PDAC cells, with various conventional RBPs exhibiting enhanced RNA association, whilst non-conventional RBPs dissociate from RNA.

Although many of the observed alterations in the RBPome are due to changes in the expression of RBPs, some RBPs show modulations in their RNA-binding activity. Specifically, a network of conventional RBPs that includes the nucleolar protein Nucleolin (Ncl), exhibits a significant enhancement in their RNA-binding activity upon oncogenic Kras induction. Using quantitative phospho-proteomics, we reveal that several of these RBPs, including Ncl, are phosphorylated downstream of Erk1/2. Phosphorylation of Ncl acts to enhance its binding to pre-ribosomal-RNA (pre-rRNA), which in turn boosts rRNA synthesis and promotes ribosome biogenesis downstream of oncogenic Kras. Crucially, we demonstrate that the enhancement of ribosome biogenesis by Ncl is essential for oncogenic Kras-induced PDAC cell proliferation and tumorigenesis, and can be targeted therapeutically to inhibit PDAC growth, *in vivo*. Our findings reveal a switch in the RNA-binding activity of Ncl, triggered by phosphorylation downstream of Erk1/2, which governs rRNA synthesis and ribosome biogenesis, and demonstrate a key targetable role for this process in RAS-mediated tumors.

## Results

### Oncogenic Kras reshapes the RNA-binding landscape of mouse PDAC cells

To study oncogenic Kras signaling, we employed tumor cells from an inducible mouse model of PDAC (iKras), in which oncogenic Kras^G12D^ expression can be controlled by administration of doxycycline (Ying *et al*, 2012). We confirmed that removal of doxycycline and the consequent loss of Kras^G12D^ expression resulted in downregulation of Erk1/2 signaling in this model (Figure S1A & S1B). Conversely, addition of doxycycline to doxycycline-withdrawn cells induced Kras^G12D^ expression and Erk1/2 activation (Figure S1C and S1D). To study the RBPome, we employed Orthogonal Organic Phase Separation (OOPS), a whole-transcriptome RIC method which uses UV-C crosslinking coupled with phenol-chloroform based phase separation to purify *in vivo* crosslinked RNA-Protein moieties (Queiroz *et al*., 2019). We validated that OOPS specifically enriched for crosslinked RBPs (Figure S1E & S1F). We then devised a quantitative RIC (qRIC) approach by combining OOPS with Stable Isotope Labelling by Amino acids in Culture (SILAC) (Ong & Mann, 2006), allowing us to measure changes in the RBPome upon induction of Kras^G12D^ expression (Figure 1A). Inducing Kras^G12D^ expression resulted in a significant increase in the RNA-bound levels of 73 proteins, whilst 101 proteins showed a significant decrease (Figure 1B & Dataset S1). Category enrichment analysis revealed that the majority of increased RNA-bound proteins were conventional RBPs, belonging to various RBP families involved in nuclear RNA processing and ribosome biogenesis (Figure 1C & Dataset S2). In contrast, the majority of decreased RNA-bound proteins were non-conventional RBPs such as metabolic enzymes and cytoskeletal proteins (Figure 1D & Dataset S3). Importantly, the effect of Kras^G12D^ on the RBPome was largely abrogated upon treating the cells with Trametinib, a specific MEK kinase inhibitor which blocks Erk1/2 activation downstream of Kras^G12D^ (Wright & McCormack, 2013), suggesting that the observed changes are largely dependent on Erk1/2 signaling (Figure 1E, S1G-H & Dataset S4).

**Figure 1:**
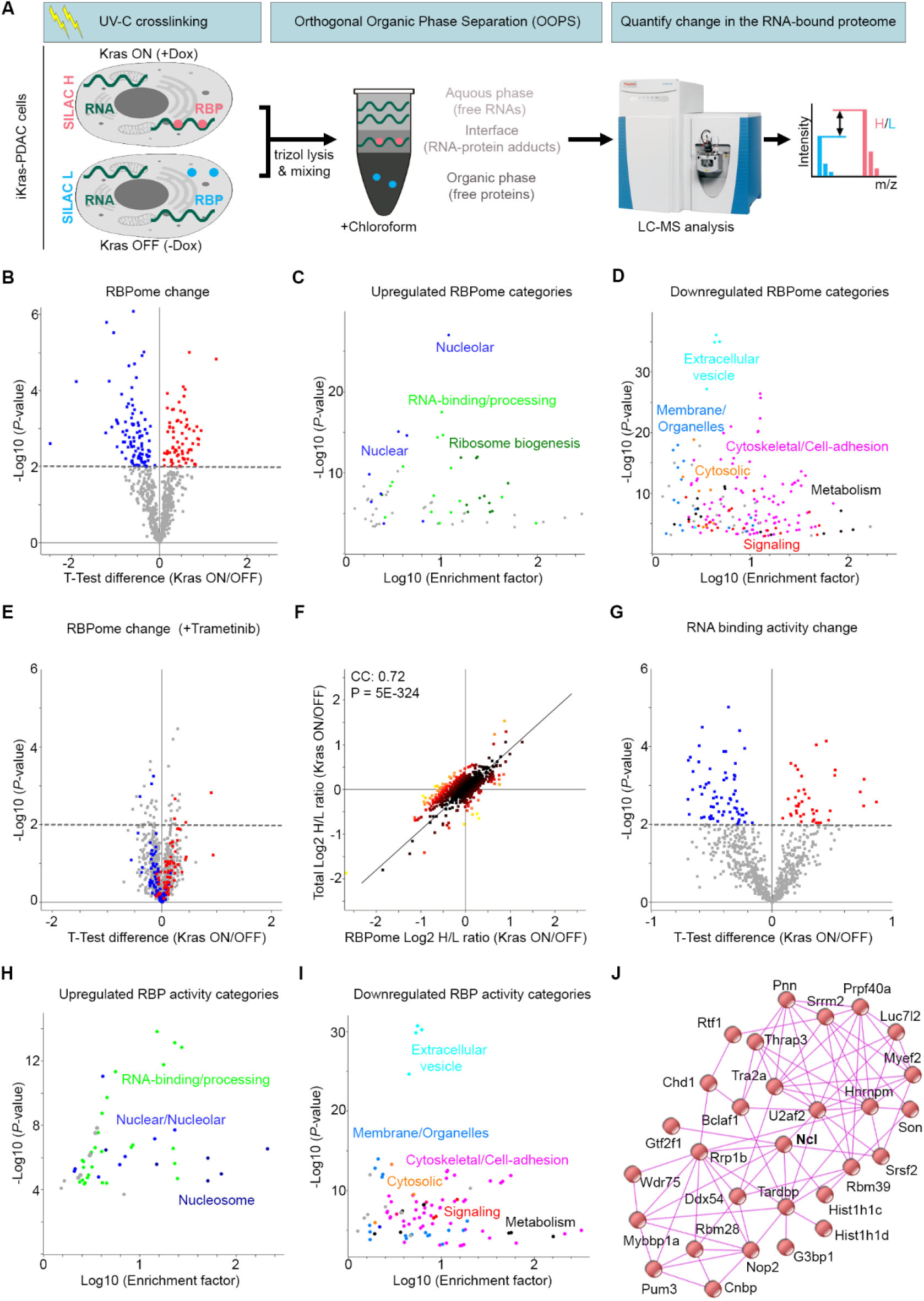
Kras^G12D^ reshapes the RBPome landscape of PDAC cells. **(A)** Experimental workflow for qRIC analysis. Heavy (H) and Light (L) SILAC labelled iKras cells, with or without doxycycline (Dox) to induce Kras^G12D^ expression, were subjected to UV-C crosslinking, TRIzol lysis, mixing of differentially labelled conditions, and OOPS analysis as in (Queiroz *et al*., 2019). RNA-crosslinked proteins, which separate into the interface, were then extracted and subjected to Trypsin digestion and liquid chromatography coupled with mass spectrometry LC-MS analysis. The SILAC (H/L) ratio values revealed Kras^G12D^-regulated changes in the RBPome. **(B)** Volcano plot of changes in the RBPome following Kras^G12D^ induction. RBPome changes were quantified, as described in (A), from six independent qRIC experiments (Dataset S1), using a one-sample t-test analysis. 73 proteins were upregulated in the RNA-bound fraction (red), whilst 101 RBPs showed a significant decrease (blue) (P < 0.01). **(C)** Fisher’s exact test analysis of protein categories that are over-represented amongst the upregulated RBPome (FDR < 0.02). Each data point represents a category from Gene Ontology (GO) and Kyoto Encyclopedia of Genes & Genome (KEGG) databases, with functionally similar categories highlighted with the same colors (Dataset S2). **(D)** Fisher’s exact test analysis of protein categories that are over-represented amongst the downregulated RBPome (FDR < 0.02). Each data point represents a category from GO and KEGG databases, with functionally similar categories highlighted with the same colors (Dataset S3). **(E)** Volcano plot of Kras^G12D^-driven changes in the RBPome in presence of Trametinib (10 nM). RBPome changes in presence of Trametinib were quantified from four independent qRIC experiments, using a one-sample t-test analysis (P < 0.01), with significantly increased (red) or decreased (blue) proteins from (B) highlighted on the plot (Dataset S4). **(F)** SILAC Total proteome changes following Kras^G12D^ induction were measured and plotted against RBPome changes from (B). Pearson’s Correlation Coefficient (CC) and significance of correlation (P) between the total proteome and the RBPome changes were calculated and displayed on the graph. **(G)** Volcano plot of changes in the RNA-binding activity following Kras^G12D^ induction. RBPome ratio changes from (B) were normalized to total proteome ratio changes from (F) to calculate changes in the RNA-binding activity (Dataset S5), using a one-sample t-test analysis. 40 proteins showed an increase in their RNA-binding activity (red), whilst 65 proteins exhibited a decrease (blue) (P < 0.01). **(H)** Fisher’s exact test analysis of categories that are over-represented amongst the proteins with enhanced RNA-binding activity (FDR < 0.02). Each data point represents a category from GO and KEGG databases, with functionally similar categories highlighted with the same colors (Dataset S6). **(I)** Fisher’s exact test analysis of categories that are over-represented amongst the proteins with diminished RNA-binding activity (FDR < 0.02). Each data point represents a category from GO and KEGG databases, with functionally similar categories highlighted with the same colors (Dataset S7). **(J)** Interaction network analysis of proteins with enhanced RNA-binding activity, using the STRING physical interactions database (Franceschini *et al*, 2013).

Next, we investigated the mechanisms of RBPome modulation in response to induction of Kras^G12D^ expression. Dysregulation of RBPs in cancer is primarily thought to be brought about by changes in their expression (Pereira *et al*, 2017). However, signaling pathways have also been shown to modulate the activity of some RBPs through post-translational modifications (Hong *et al*, 2017; Matter *et al*, 2002; Tripathi *et al*, 2010). To assess these two possibilities, we performed SILAC based total proteomics on the iKras PDAC cells, with or without the induction of Kras^G12D^ expression. We observed a strong overall correlation between changes in the RBPome and the total proteome, suggesting that the majority of variation in the RBPome is likely reflective of changes in the expression levels of RBPs (Figure 1F). However, not all changes were correlative, so to specifically reveal alterations in the RBPome that were independent of protein expression modulations, we normalized the RBPome SILAC ratio values to those of the total proteome. Analysis of this RBPome to total normalized ratio value revealed changes in the RNA-binding activities of RBPs in response to Kras^G12D^ induction. We observed a significant increase in the RNA-binding activity of 40 proteins, coupled with a significant decrease in the activity of 65 proteins (Figure 1G & Dataset S5). Similar to the RBPome analysis, category enrichment revealed that the majority of proteins with increased RNA-binding activity were conventional RBPs belonging to various nuclear protein families (Figure 1H & Dataset S6), whilst the majority of proteins with decreased RNA-binding activity were non-conventional RBPs belonging mostly to metabolic and cytoskeletal protein families (Figure 1I & Dataset S7). Interactome analysis of the RBPs with increased RNA-binding activity revealed that many of them are known to physically associate with one another (Figure 1J). Several of these RBPs such as Ncl, Rrp1b, Wdr75, Nop2, and Mybbp1a are known regulators of ribosome biogenesis (Chamousset *et al*, 2010; de Beus *et al*, 1994; Liu *et al*, 2017; Moudry *et al*, 2021; Ugrinova *et al*, 2018). Together, these results reveal that Kras^G12D^ mediated Erk1/2 activation reshapes the RBPome landscape of PDAC cells, with a network of nuclear RBPs, which includes Ncl and several other ribosome biogenesis factors, exhibiting a significant enhancement in their RNA-binding activity.

### Oncogenic Kras signaling enhances the RNA-binding activity of Ncl through phosphorylation

We next set out to investigate the mechanism by which Erk1/2 signaling modulated the RNA-binding activity of RBPs downstream of Kras^G12D^. Erk1/2 is a major cellular kinase that can directly or indirectly phosphorylate a plethora of downstream cellular substrates (Yoon & Seger, 2006). We therefore hypothesized that some of the changes in RNA-binding activity could be mediated via Erk1/2 dependent phosphorylation of RBPs. To reveal Erk1/2-dependent phosphorylation changes that are brought about by Kras^G12D^ induction, we carried out a multi-variate quantitative phospho-proteomics analysis of iKras PDAC cells, using Tandem Mass Tagging (TMT) (McAlister *et al*, 2012) (Dataset S8). 7,478 phosphorylations were identified, 872 of which were found to be significantly increased upon Kras^G12D^ induction (Figure 2A). As expected, phospho-motif analysis revealed the ERK substrate motif to be highly over-represented amongst these phosphorylation sites, but motifs for a number of other kinases such as Casein Kinase I (CK1) and Casein Kinase II (CK2) were also significantly over-represented, albeit to a lesser extent (Figure S2A). Several RBPs whose RNA-binding activity was significantly enhanced upon Kras^G12D^ induction were amongst the phosphorylation targets of Kras^G12D^, often found to be phosphorylated on multiple residues (Figure 2A & 2B). The impact of Kras^G12D^ induction on the phospho-proteome was abrogated by treating the cells with Trametinib (Figure 2C), suggesting that Erk1/2 signaling is the principal driver of phosphorylation events downstream of Kras^G12D^ in iKras PDAC cells. These findings are in agreement with previous studies which showed cell-autonomous Kras^G12D^ signaling in PDAC to be primarily mediated via Erk1/2 (Tape *et al*, 2016), and reveal that a number of RBPs, whose RNA-binding activity is enhanced downstream of Kras^G12D^, undergo phosphorylation downstream of Erk1/2.

**Figure 2:**
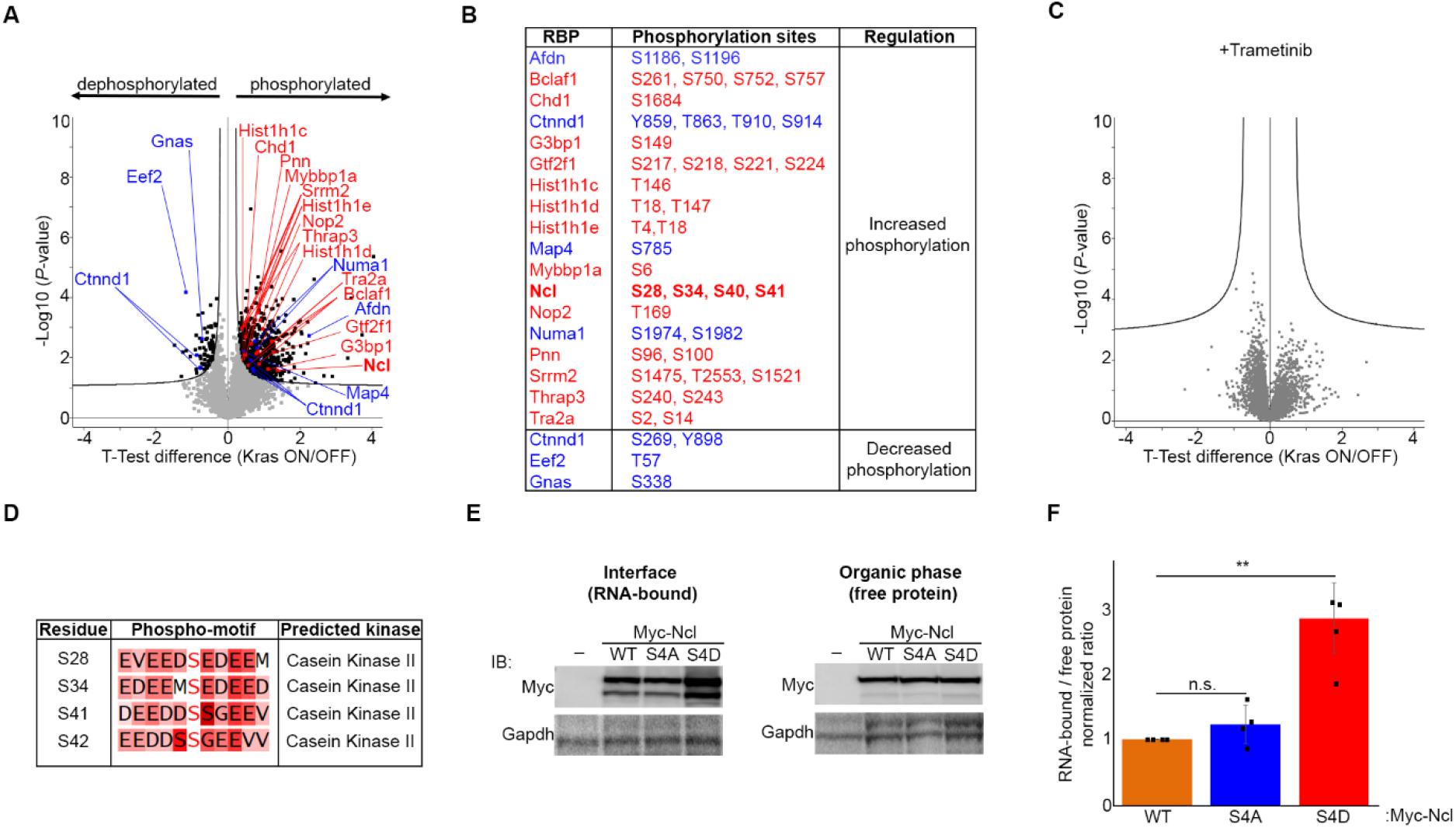
The RNA-binding activity of Ncl is enhanced upon its phosphorylation downstream of Kras^G12D^-induced Erk1/2 signaling. **(A)** Volcano plot of phosphorylation changes in iKras PDAC cells in response to Kras^G12D^ induction. Phospho-to total proteomic changes were quantified from three independent replicates of Dox treated vs. untreated cells, via TMT (Dataset S8). 872 phosphorylations were found to be significantly increased, while 82 phosphorylations were significantly decreased (FDR < 0.05). Significantly changing phosphorylations on RBPs whose RNA-binding activity was significantly increased (red) or decreased (blue) as per Figure 1G are highlighted on the plot. **(B)** List of significantly increased or decreased phosphorylation sites on the RBPs whose RNA-binding activity was enhanced (red) or reduced (blue) upon Kras^G12D^ induction. **(C)** Volcano plot of phosphorylation changes in iKras PDAC cells in response to Kras^G12D^ induction in presence of Trametinib (10 nM). Phospho-to total proteomic changes were quantified from three independent replicates of Dox + Trametinib treated vs. untreated cells (Dataset S8). No phosphorylations were found to be significantly changing upon Kras^G12D^ induction in presence of Trametinib (FDR < 0.05). **(D)** Phospho-motif kinase prediction analysis of Kras^G12D^-induced Ncl phosphorylation sites, using NetworKIN (Linding *et al*, 2008). **(E)** OOPS analysis of the RNA-binding activity of WT, Phospho-defective (S4A), and phospho-mimicking (S4D) mutants of Ncl. Vectors encoding Myc-tagged WT, S4A, and S4D Ncl were transiently expressed in iKras PDAC cells, before OOPS and western blot analysis of the interface (RNA-bound) and the organic phase (free protein) fractions with anti-Myc antibody. Gapdh, which also binds RNA (Dataset S1), was blotted as loading control. **(E)** Quantification of normalized RNA-bound to free Myc-Ncl ratio values from (E), as measures of RNA-binding activity (n = 4) (**: P < 0.01; n.s.: not significant).

Amongst the Kras^G12D^ regulated RBPs which were significantly phosphorylated downstream of Erk1/2, Ncl was strongly phosphorylated on four closely situated Serine residues (S28, S34, S40, and S41) within its N-terminal region (Figure 2B). This region, consisted of acidic-rich regions interspersed with stretches of basic residues, is predicted to be largely disordered (Jumper *et al*, 2021). It is also well-known to undergo phosphorylation by multiple kinases (Tajrishi *et al*, 2011), but no direct link between Ncl phosphorylation and Erk1/2 signaling has been reported. Phospho-motif analysis revealed that the identified phosphorylation sites are not direct Erk1/2 targets, but most likely phosphorylated by CK2 (Figure 2D), which is well-known to physically associate with and phosphorylate Ncl (Caizergues-Ferrer *et al*, 1987; Li *et al*, 1996). Importantly, CK2 has been shown to be activated downstream of ERK1/2 signaling (Ji *et al*, 2009), which is in agreement with our phospho-motif analysis of the Kras^G12D^ enhanced phosphorylation sites (Figure S2A). Accordingly, we could demonstrate by using a specific phospho-CK2-Substrate antibody mix that Kras^G12D^ induction enhanced CK2 activity in an Erk1/2-dependent manner in iKras PDAC cells (Figure S2B-E). These findings suggest that Erk1/2 activation downstream of Kras^G12D^ increases CK2 activity, leading to enhanced phosphorylation of Ncl on S28, S34, S40, and S41.

To reveal whether the observed phosphorylations and the enhancement of the RNA-binding activity of Ncl were causally linked, we generated phospho-defective and phospho-mimicking mutants of Ncl, in which the four identified residues were mutated to either Alanine (S4A), or Aspartic acid (S4D), respectively. Myc-tagged versions of these mutants along with the wild-type (WT) control were ectopically expressed in iKras PDAC cells, and the cells were subjected to OOPS analysis. The interface, which contains RNA-bound proteins, along with the organic phase, which contains free proteins, were then analyzed by western blotting with an anti-Myc antibody. While no variations were observed between the levels of Ncl in the organic phase, the levels of the phospho-mimicking (S4D) mutant exhibited a significant increase in the interface, suggesting that its RNA-binding activity must be significantly increased (Figure 2E & 2F). These results suggest that phosphorylation of Ncl on S28, S34, S40, and S41 acts to enhance its RNA-binding activity downstream of Kras^G12D^.

### Transcriptome-wide iCLIP studies reveal Ncl to be primarily associated with pre-rRNA

After revealing that the RNA-binding activity of Ncl is enhanced by phosphorylation downstream of Kras^G12D^, we set out to determine the full spectrum of RNAs that are bound by Ncl and their dynamics in response to Kras^G12D^-dependent phosphorylation. Although predominantly known to be localized to the nucleolus, where it regulates several steps of ribosome biogenesis, Ncl has also been shown to have diverse extra-nucleolar functions in the nucleoplasm, cytoplasm, as well as the cell-surface (Ugrinova *et al*., 2018). These include regulation of chromatin architecture (Angelov *et al*, 2006; Erard *et al*, 1988), microRNA processing (Pichiorri *et al*, 2013; Pickering *et al*, 2011), translation (Abdelmohsen *et al*, 2011; Takagi *et al*, 2005), as well as mRNA turnover (Sengupta *et al*, 2004; Zhang *et al*, 2006). In addition, cell-surface localized Ncl has been shown to be involved in regulation of cell adhesion and transmembrane signaling (Losfeld *et al*, 2009; Reyes-Reyes & Akiyama, 2008). In order to identify the full repertoire of RNAs that directly interact with Ncl, and reveal the impact of Kras^G12D^-dependent phosphorylation on their association with Ncl, we performed individual-nucleotide resolution UV crosslinking and immunoprecipitation (iCLIP) (Konig et al., 2010). Ncl-RNA complexes were purified by immunoprecipitation with an anti-Myc-tag antibody, from iKras PDAC cells that ectopically expressed Myc-tagged WT, S4A, or S4D mutants of Ncl, or a mock transfected negative control. Western blot analysis revealed that the ectopic proteins were expressed at around 50% of the endogenous levels, ruling out potential artefacts due to high levels of overexpression (Figure S3A & S3B). Moreover, little mapped iCLIP reads were identified from the negative control iCLIP samples, as opposed to the Myc-Ncl samples, suggesting that the identification of Ncl-interacting RNAs was highly specific (Figure S3C).

More than 70% of all the Myc-Ncl iCLIP mapped reads corresponded to the ribosomal-DNA (rDNA) locus, which codes for a long primary transcript known as the 47S pre-rRNA that is ultimately processed into 18S, 5.8S, and 28S rRNAs (Figure 3A). No significant difference between WT and the phospho-mutants was observed, suggesting that phospho-regulation of Ncl by Kras^G12D^ does not affect its repertoire of RNA substrates (Figure 3A). In agreement with these results, immunofluorescence analysis revealed that ectopically expressed Myc-tagged WT, S4A, and S4D mutants of Ncl were exclusively localized to the nucleolus (Figure 3B). Localization of ectopic Myc-Ncl was in complete accordance with endogenous Ncl, which was also found to be exclusively nucleolar, irrespective of Kras^G12D^ status (Figure S3D). Together, these results suggest that Ncl is primarily localized to the nucleolus of iKras PDAC cells where it predominantly binds pre-rRNA, irrespective of its phosphorylation status.

**Figure 3:**
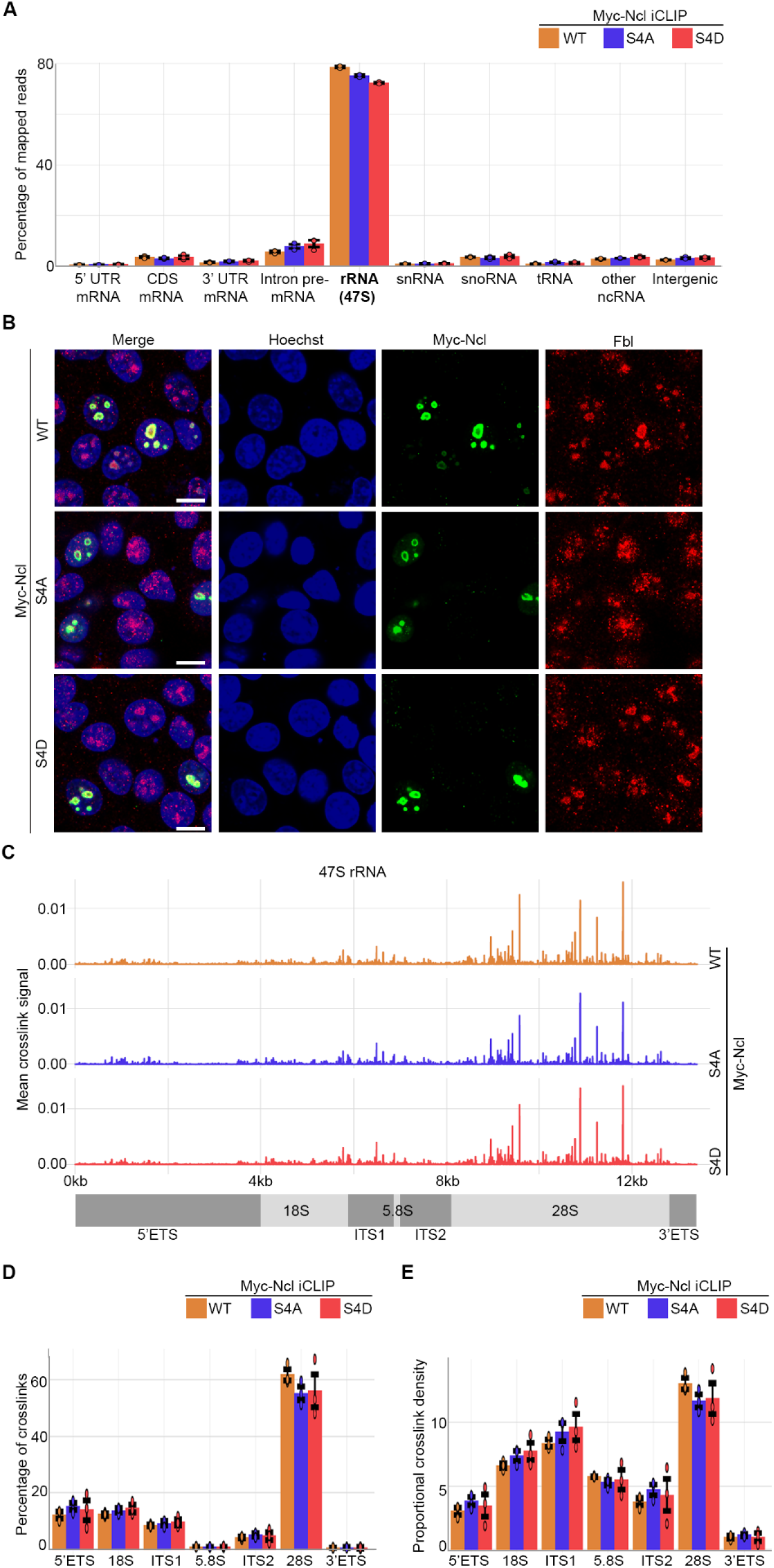
Ncl is predominantly bound to pre-rRNA, irrespective of its phosphorylation status. **(A)** Percentage of mapped reads belonging to different types of RNA from three independent iCLIP replicate experiments of WT, S4A, and S4D Myc-Ncl. **(B)** Immunofluorescence analysis of WT, S4A, and S4D Myc-Ncl subcellular localization. IKras PDAC cells ectopically expressing either WT, S4A, or S4D Myc-Ncl were fixed and immunostained with anti-Myc-tag antibody, along with anti-Fibrillarin (Fbl) antibody as a Nucleolar marker, and Hoechst, followed by confocal microscopy analysis. Scale bar = 10 μm. **(C)** Distribution of WT, S4A, and S4D Myc-Ncl crosslink sites across the annotated 47S pre-rRNA genomic region. Peak heights represent mean crosslink intensities from three independent iCLIP replicate experiments. **(D)** Quantification of the percentage of crosslinks in each region of the 47S pre-rRNA for WT, S4A, and S4D Myc-Ncl. Percentage of crosslinks relative to 47S total were quantified from three independent iCLIP experiments. **(E)** Quantification of the proportional density of crosslinks in each region of the 47S pre-rRNA for WT, S4A, and S4D Myc-Ncl. Proportional density was calculated from three independent iCLIP experiments by normalizing the number of crosslinks in each region to the sequence length of that region.

Although Kras^G12D^-dependent phosphorylation of Ncl does not affect its primary association with pre-rRNA, the enhancement of the RNA-binding activity could be resulting in generation of novel Ncl binding sites on the pre-rRNA. Alternatively, Ncl could be binding to exactly the same sites, but with higher affinity. To differentiate between these possibilities, we analyzed the distribution of Ncl crosslink sites within the pre-rRNA. 47S pre-rRNA contains the sequences of 18S, 5.8S, and 28S rRNAs, flanked by two external transcribed spacers at each end (5’ETS and 3’ETS), as well as two Internal transcribed spacers (ITS1 and ITS2) that are situated between 18S, 5.8S, and 28S. During rRNA processing, these spacer sequences are removed in a step-by-step manner via the action of different endo- and exo-ribonucleases (Turowski & Tollervey, 2015). WT Ncl showed binding throughout the length of 47S pre-rRNA, with numerous crosslink spikes detected in various regions (Figure 3C). No difference between the crosslinking patterns of the WT and the phospho-mutants were observed, suggesting that phosphorylations do not change the binding pattern of Ncl on pre-rRNA (Figure 3C). Accordingly, quantification of the percentage as well as the density of Ncl crosslinks within each pre-rRNA region revealed no differences between WT Ncl and the phospho-mutants (Figure 3D & 3E). Collectively, these results suggest that the Kras^G12D^-dependent phosphorylations of Ncl do not affect its pattern of binding to pre-rRNA. We therefore conclude that the Kras^G12D^-dependent phospho-regulation of Ncl must be acting to enhance its RNA-binding affinity, without affecting its RNA-binding specificity.

### Ncl phosphorylation enhances rRNA synthesis and ribosome biogenesis

The predominant interaction of Ncl with pre-rRNA suggests that it must be primarily functioning in regulating ribosome biogenesis in iKras PDAC cells, so we next investigated the functional impact of Ncl phospho-regulation on controlling ribosome biogenesis downstream of Kras^G12D^. We first assessed the impact of Kras^G12D^ induction on nascent rRNA levels, using single-cell visualization of newly synthesized RNAs by 5-fluorouridine (FUrd) pulse-labeling (Percipalle & Louvet, 2012). Induction of Kras^G12D^ expression by addition of doxycycline to doxycycline-withdrawn cells triggered a strong accumulation of nascent RNA within the nucleolus of iKras PDAC cells. This accumulation was largely abrogated by short-term treatment of the cells with an rRNA synthesis inhibitor (CX-5461), suggesting that the nucleolar nascent RNA signal must be largely comprised of newly synthesized rRNA (Figure S4A & S4B). Conversely, removal of doxycycline and the consequent loss of Kras^G12D^ expression resulted in a substantial reduction in nascent rRNA levels (Figure S4C & S4D), collectively demonstrating a strong dependence of rRNA synthesis on Kras^G12D^ expression. Crucially, depletion of Ncl via two independent siRNA oligos abrogated nascent rRNA accumulation downstream of Kras^G12D^ (Figure 4A, 4B, & S4E). As an independent approach, we also measured the levels of unprocessed pre-rRNA by quantitative reverse transcription-PCR (RT-qPCR), using a probe against the 5’ETS region of 47S pre-rRNA which is removed at the first step of pre-rRNA processing (Pineiro *et al*, 2018). In agreement with the nascent RNA imaging results, Kras^G12D^ expression resulted in an increase in the levels of unprocessed pre-rRNA, but this increase was abrogated upon Ncl depletion (Figure 4C). Together, these findings suggest that Kras^G12D^ acts to enhance pre-rRNA expression via Ncl.

**Figure 4:**
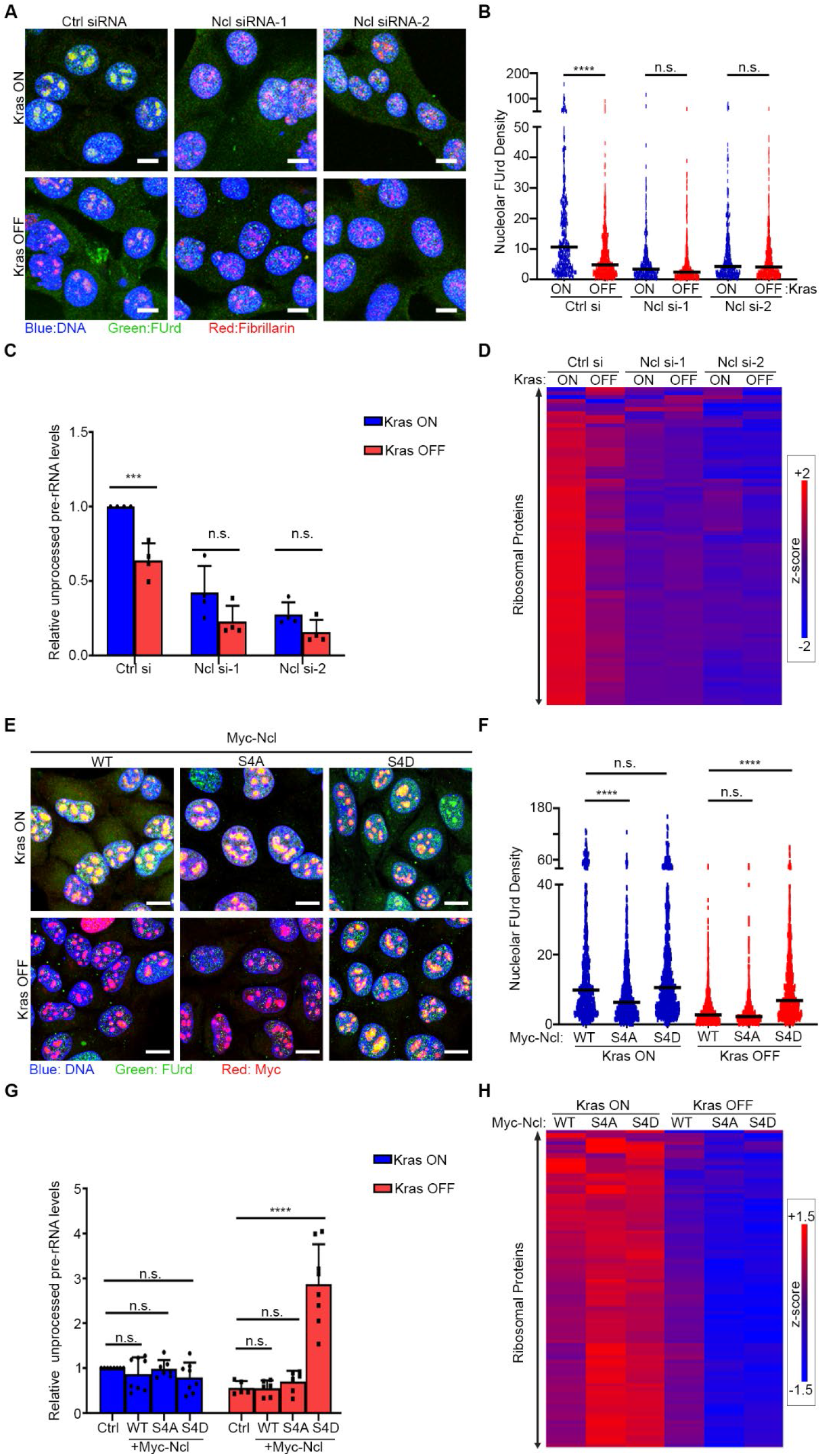
Kras^G12D^ promotes rRNA synthesis and ribosome biogenesis via Ncl. **(A)** Nascent RNA imaging in control and Ncl depleted iKras PDAC cells, in presence or absence of Kras^G12D^. Cells were transfected with a non-targeting control siRNA, or two independent siRNAs against Ncl, grown for 48hrs in presence or absence of Dox, before being subjected to pulse labeling with FUrd. Cells were then fixed and immunostained with anti-FUrd antibody (green) to visualize nascent RNA, along with anti-Fibrillarin (Fbl) antibody as a Nucleolar marker (red), and Hoechst (blue) as the Nuclear stain, followed by confocal microscopy analysis. Scale bar = 10 μm. **(B)** Quantification of Nucleolar FUrd levels in images from (A). FUrd fluorescence densities in single nucleoli were quantified from 182-220 individual cells per condition, combined from two independent biological replicate experiments (****: P < 0.0001; n.s.: not significant). **(C)** RT-qPCR analysis of 5’ETS-containing pre-rRNA transcript levels in control and Ncl depleted iKras PDAC cells, in presence or absence of Kras^G12D^. Cells were transfected with a non-targeting control siRNA, or two independent siRNAs against Ncl, and grown for 48hrs in presence or absence of Dox, before RT-qPCR analysis with a specific probe against the mouse 5’ETS region. A probe against mouse Actb mRNA was used as loading control for normalization (n = 3) (***: P < 0.001; n.s.: not significant). **(D)** TMT quantitative analysis of RP levels in control and Ncl depleted iKras PDAC cells, in presence or absence of Kras^G12D^. Cells were transfected with a non-targeting control siRNA, or two independent siRNAs against Ncl, and grown for 48hrs in presence or absence of Dox, before lysis and TMT-mediated quantitative proteomics analysis. Z-scores of TMT intensity changes for all identified RPs across the different conditions were plotted as a heat map (red → increase; blue → decrease). **(E)** Nascent RNA imaging of WT, S4A, and S4D Myc-Ncl expressing iKras PDAC cells, in presence or absence of Kras^G12D^. Vectors encoding Myc-tagged WT, S4A, and S4D Ncl were transiently transfected into iKras PDAC cells, before reseeding and growing the cells for 48hrs in presence or absence of Dox. Cells were then subjected to pulse labeling with FUrd, fixation, and immunostaining with anti-FUrd antibody (green), anti-Myc-tag antibody (red), and Hoechst (blue), followed by confocal microscopy analysis. Scale bar = 10 μm. **(F)** Quantification of FUrd levels in Myc-positive nucleoli from (E). FUrd fluorescence densities in single nucleoli were quantified from 160-281 individual cells per condition, combined from three independent biological replicate experiments (****: P < 0.0001; n.s.: not significant). **(G)** RT-qPCR analysis of 5’ETS-containing pre-rRNA transcript levels in WT, S4A, and S4D Myc-Ncl expressing iKras PDAC cells, in presence or absence of Kras^G12D^. Vectors encoding Myc-tagged WT, S4A, and S4D Ncl were transiently transfected into iKras PDAC cells, before reseeding and growing the cells for 48hrs in presence or absence of Dox, followed by RT-qPCR analysis with a specific probe against the mouse 5’ETS region. A probe against mouse Actb mRNA was used as loading control for normalization (n = 6) (****: P < 0.0001; n.s.: not significant). **(H)** TMT quantitative analysis of RP levels in WT, S4A, and S4D Myc-Ncl expressing iKras PDAC cells, in presence or absence of Kras^G12D^. Vectors encoding Myc-tagged WT, S4A, and S4D Ncl were transiently transfected into iKras PDAC cells, before reseeding and growing the cells for 48hrs in presence or absence of Dox. Cells were then lysed and analyzed by TMT-mediated quantitative proteomics. Z-scores of TMT intensity changes for all identified RPs across the different conditions were plotted as a heat map (red → increase; blue → decrease).

Synthesis of 47S pre-rRNA is a critical step in the regulation of eukaryotic ribosome biogenesis (Pelletier *et al*, 2018), so we next assessed whether the Ncl-dependent enhancement of pre-rRNA levels by Kras^G12D^ leads to an increase in cellular ribosome levels. Time-course TMT quantitative proteomics analysis of iKras PDAC cells revealed a significant accumulation of Ribosomal Proteins (RPs) in response to induction of Kras^G12D^ expression (Figure S4F & Dataset S9). Conversely, RPs were depleted over time in response to loss of Kras^G12D^ expression (Figure S4G & Dataset S10). In fact, category enrichment analysis revealed that protein categories corresponding to RPs and rRNA processing were amongst the most depleted in response to Kras^G12D^ loss (Figure S4H & Dataset S11). Since RPs are known to be highly unstable unless incorporated into mature ribosomal subunits (Lam *et al*, 2007), their stable accumulation in response to Kras^G12D^ expression is indicative of more ribosomes having been synthesized. Crucially, Ncl depletion abrogated the stable accumulation of RPs in response to Kras^G12D^ expression, suggesting that enhancement of ribosome biogenesis downstream of Kras^G12D^ is dependent on Ncl (Figure 4D).

To determine whether the Erk1/2-dependent phosphorylation of Ncl plays a role in regulating pre-rRNA expression and ribosome biogenesis downstream of Kras^G12D^, we evaluated the impact of phospho-defective (S4A) and phospho-mimicking (S4D) mutants of Ncl on nascent pre-rRNA levels, in presence or absence of Kras^G12D^. In presence of Kras^G12D^, iKras PDAC cells ectopically expressing either WT Ncl or the phospho-mutants exhibited accumulation of nascent rRNA in their nucleolus, though nascent rRNA levels were mildly but significantly decreased in S4A expressing cells (Figure 4E & 4F). Kras^G12D^ removal abrogated nascent rRNA levels in the WT and S4A mutant expressing cells, but the expression of the S4D mutant could rescue nascent rRNA expression in the absence of Kras^G12D^ (Figure 4E & 4F), suggesting that phosphorylation of Ncl is sufficient for enhancing nascent rRNA expression downstream of Kras^G12D^. RT-qPCR analysis revealed that the S4D mutant caused a drastic accumulation of unprocessed pre-rRNA in the absence of Kras^G12D^ (Figure 4G). This accumulation far exceeded the pre-rRNA levels in presence of Kras^G12D^, suggestive of a defect in pre-rRNA processing (Dermit *et al*., 2020). Based on these results, we conclude that the phospho-mimicking S4D mutant can enhance nascent pre-rRNA expression in the absence of Kras^G12D^, but the cells are defective in mediating the processing of pre-rRNA. Accordingly, quantitative proteomics analysis of iKras PDAC cells that ectopically expressed WT Ncl or the phospho-mutants revealed no S4D mediated rescue of ribosome biogenesis in the absence of Kras^G12D^ expression (Figure 4H). Together, these findings suggest that Ncl phosphorylation downstream of Kras^G12D^ acts to enhance pre-rRNA expression, but the subsequent processing of nascent pre-rRNA likely requires other KRAS^G12D^-activated factors.

### Ncl is crucial for Oncogenic Kras mediated PDAC cell proliferation and tumorigenesis

Hyperactive ribosome biogenesis is a hallmark of most malignancies, acting to sustain augmented protein synthesis that underpins unrestricted cancer cell proliferation and tumor growth (Pelletier *et al*., 2018). We therefore investigated whether the Ncl-mediated enhancement of ribosome biogenesis downstream of Kras^G12D^ was crucial for PDAC proliferation and tumor formation. As demonstrated before (Ying *et al*., 2012), Kras^G12D^ removal significantly reduced the proliferation of iKras PDAC cells in long-term clonogenic assays under standard 2D cell culture settings (Figure S5A, & S5B). The impact of Kras^G12D^ expression on PDAC cell proliferation was also evident in a 3D matrix made up of collagen-I, which forms the bulk of PDAC extracellular matrix, with rapid proliferative growth of the cell mass into the matrix that was strictly dependent on Kras^G12D^ expression (Figure S5C & S5D). Depletion of Ncl abrogated the effect of Kras^G12D^ in both 2D and 3D culture settings, revealing a strong dependence on Ncl for Kras^G12D^ mediated PDAC cell proliferation (Figure 5A-D).

**Figure 5:**
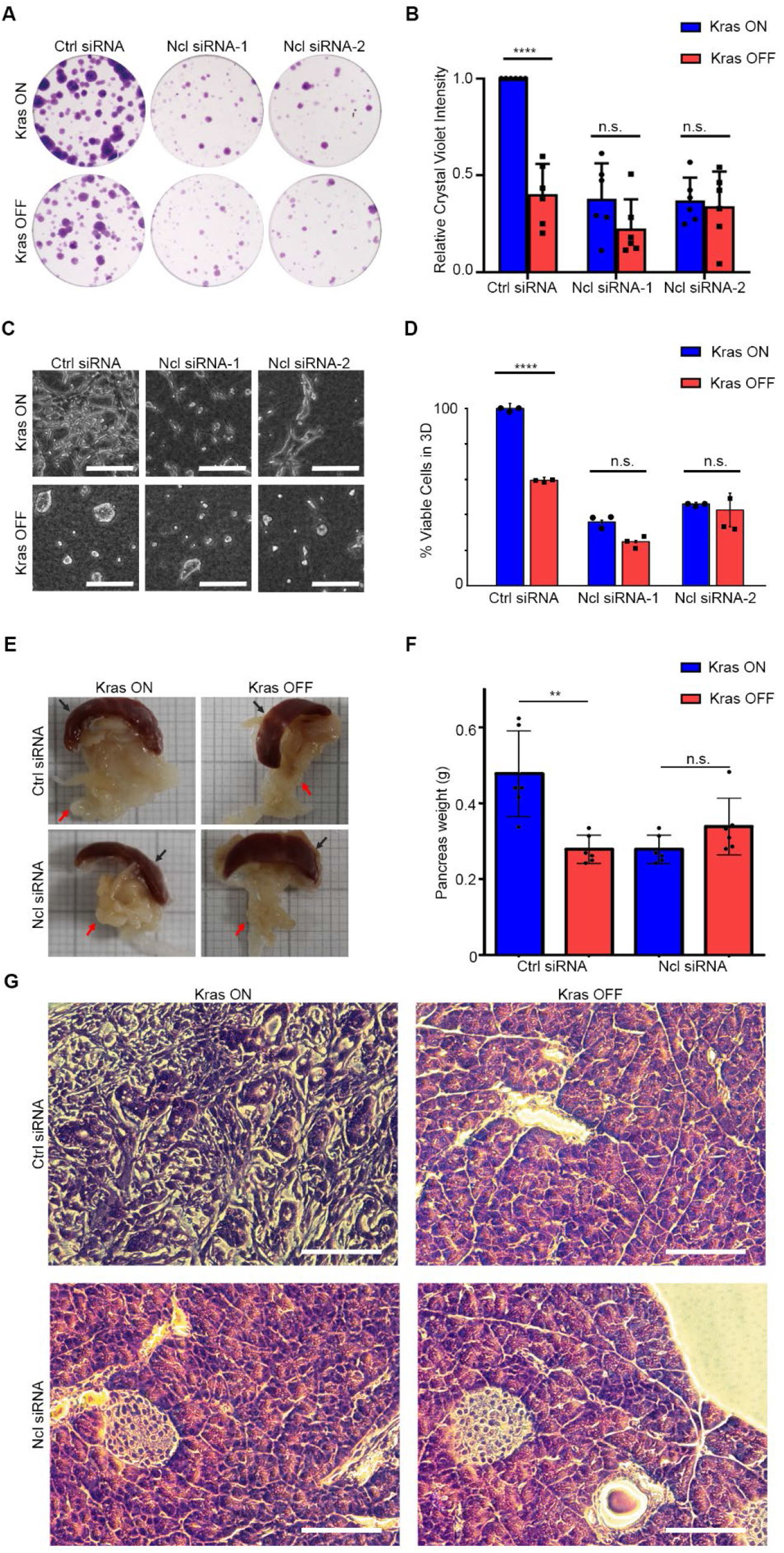
Ncl is necessary for Kras^G12D^-mediated PDAC cell proliferation and tumor formation. **(A)** Colony formation of control and Ncl depleted iKras PDAC cells, in presence or absence of Kras^G12D^. Cells were transfected with a non-targeting control siRNA, or two independent siRNAs against Ncl, followed by clonogenic assay for 7 days in presence or absence of Dox. Colonies were visualized by Crystal Violet staining. **(B)** Quantification of Crystal Violet staining levels from (A) (n = 6). (****: P < 0.0001; n.s.: not significant). **(C)** 3D proliferation of control and Ncl depleted iKras PDAC cells, in presence or absence of Kras^G12D^. Cells were transfected with a non-targeting control siRNA, or two independent siRNAs against Ncl, before being reseeded onto 3D Collagen-I gels, with or without Dox, and allowed to grow for 48 hrs. Cells were subsequently imaged live by phase contrast microscopy. Scale bar = 200 μm. **(D)** Analysis of the relative percentage of viable cells in 3D collagen-I cultures from (C). Cells were subjected to luminescence-based viability assay by CellTiter-Glo to quantify the percentage of viable cells (n = 3) (****: P < 0.0001; n.s.: not significant). **(E)** Representative images of the pancreas from control or Ncl depleted iKras PDAC engrafted mice, in presence or absence of Kras^G12D^. Non-targeting control or Ncl siRNA transfected iKras PDAC cells were orthotopically engrafted into the pancreas of nude mice. Mice were fed either Dox -containing (Kras ON) or Dox -free (Kras OFF) water for 7 days, before culling and extraction of their pancreas (red arrow). Spleen (black arrow), which is located adjacent to the pancreas, was also extracted and included in the images for comparison. **(F)** Quantification of pancreas weights from (E), as a measure of orthotopic tumor growth. Pancreas weights from six animals per condition were quantified. (**: P < 0.01; n.s.: not significant). **(G)** H&E analysis of the extracted pancreas tissues from (E). Extensive portions of the tissue in control-engrafted mice fed with Dox display PDAC histology, with malignant ductal structures surrounded by stroma. However, typical pancreas histology comprised of acini, islets, and normal ducts is observed in all other conditions. Scale bar = 100 μm.

Using orthotopic xenografts of the iKras PDAC cells, we next investigated whether Kras^G12D^-induced tumor formation *in vivo* was dependent on Ncl. Orthotopic xenografts of iKras PDAC cells have been demonstrated to generate tumors which faithfully recapitulate the histological and molecular features of PDAC, in a doxycycline-dependent manner (Ying *et al*., 2012). In our pilot studies, tumors were fully established within a week of engraftment in doxycycline-fed mice, with animals having to be sacrificed 2-3 weeks post-engraftment due to rapid disease progression (Figure S5E). Thus, we chose a time-scale of one week post-engraftment for investigating the impact of Ncl depletion on Kras^G12D^-dependent tumor formation. As expected, doxycycline-mediated Kras^G12D^ expression triggered significant tumor growth in the pancreas of the engrafted mice, which was inhibited in the absence of doxycycline. Depletion of Ncl abrogated this Kras^G12D^-induced tumor formation (Figure 5E, 5F, & S5F). Histological analysis further revealed an extensive Kras^G12D^-dependent growth of the malignant component and its associated stroma within the pancreas of the engrafted mice, which was completely inhibited upon Ncl depletion (Figure 5G). Collectively, these results reveal that Ncl is crucial for PDAC cell proliferation and tumor formation downstream of Kras^G12D^.

### Kras dependency on Ncl-mediated ribosome biogenesis can be therapeutically targeted

Our results reveal a dependency for Ncl in Kras^G12D^-mediated PDAC tumor formation, suggesting a possibility for therapeutic exploitation. Ncl has been a subject of significant interest as a therapeutic target for a diverse range of cancers, owing to its often strong upregulation of expression combined with its pro-malignancy functions (Abdelmohsen & Gorospe, 2012). However, most efforts so far have been focused on cell-surface localized Ncl, which is commonly observed in cancerous but not normal tissues. Accordingly, several compounds that can target extracellularly localized Ncl have been developed (Abdelmohsen & Gorospe, 2012), but these compounds are unlikely to reach nucleolar Ncl in significant quantities due to lack of cell-permeability. We demonstrated that Ncl is exclusively localized to the nucleolus, associated with pre-rRNA, and functions to promote ribosome biogenesis in iKras PDAC cells. We thereby reasoned that pharmacological inhibition of ribosome biogenesis should functionally mimic Ncl removal. For this purpose, we utilized CX-5461, as it can be orally administered, *in vivo* (Drygin *et al*, 2011). CX-5461 has shown promising results in early clinical trials against a number of human malignancies (Hilton *et al*, 2018; Khot *et al*, 2019). However, its *in vivo* mechanism of action has been subject to some controversy. Early reports suggested that the primary anti-tumor activity of CX-5461 arises from inhibition of ribosome biogenesis (Bywater *et al*, 2012; Drygin *et al*., 2011). However, recent studies have revealed that CX-5461 can also induce DNA damage, which seems to act as the primary cause of its cytotoxicity in several cell-lines (Bruno *et al*, 2020; Negi & Brown, 2015; Pan *et al*, 2021; Quin *et al*, 2016; Sanij *et al*, 2020; Xu *et al*, 2017). This is proposed to be initiated by the irreversible arrest of RNA polymerase I on rDNA promoter regions, leading to nucleolar stress that propagates into a genome-wide DNA damage response (Mars *et al*, 2020). Importantly, a recent study has shown that the dosage of CX-5461 could be adjusted to minimize DNA damage induction, while still achieving an effective inhibition of rRNA synthesis (Prakash *et al*, 2019). Therefore, we performed a dose-titration of CX-5461 in iKras PDAC cells, and assessed the treatment impacts on nascent rRNA expression as well as DNA damage. Short-term treatment of iKras PDAC cells with low nanomolar doses of CX-5461 induced a potent inhibition of nascent rRNA expression (Figure 6A & 6B). This was followed by a significant depletion in the levels of 47S pre-rRNA (Figure 6C). CX-5461 treatment also induced DNA damage in iKras PDAC cells, but this only occurred at the 1 μM dose (Figure S6A & S6B). Consistent with the proposed role of nucleolar stress in this process, the 1 μM dose also caused the disruption of nucleoli, as evidenced by the leakage of Ncl into the nucleoplasm, which was not observed with lower CX-5461 doses (Figure 6A). Critically, low nanomolar doses of CX-5461 could still abrogate the impact of Kras^G12D^ on the proliferation of iKras PDAC cells, suggesting that inhibition of rRNA synthesis is the critical target of CX-5461 in the context of Kras^G12D^-driven cell proliferation (Figure 6D & 6E).

**Figure 6:**
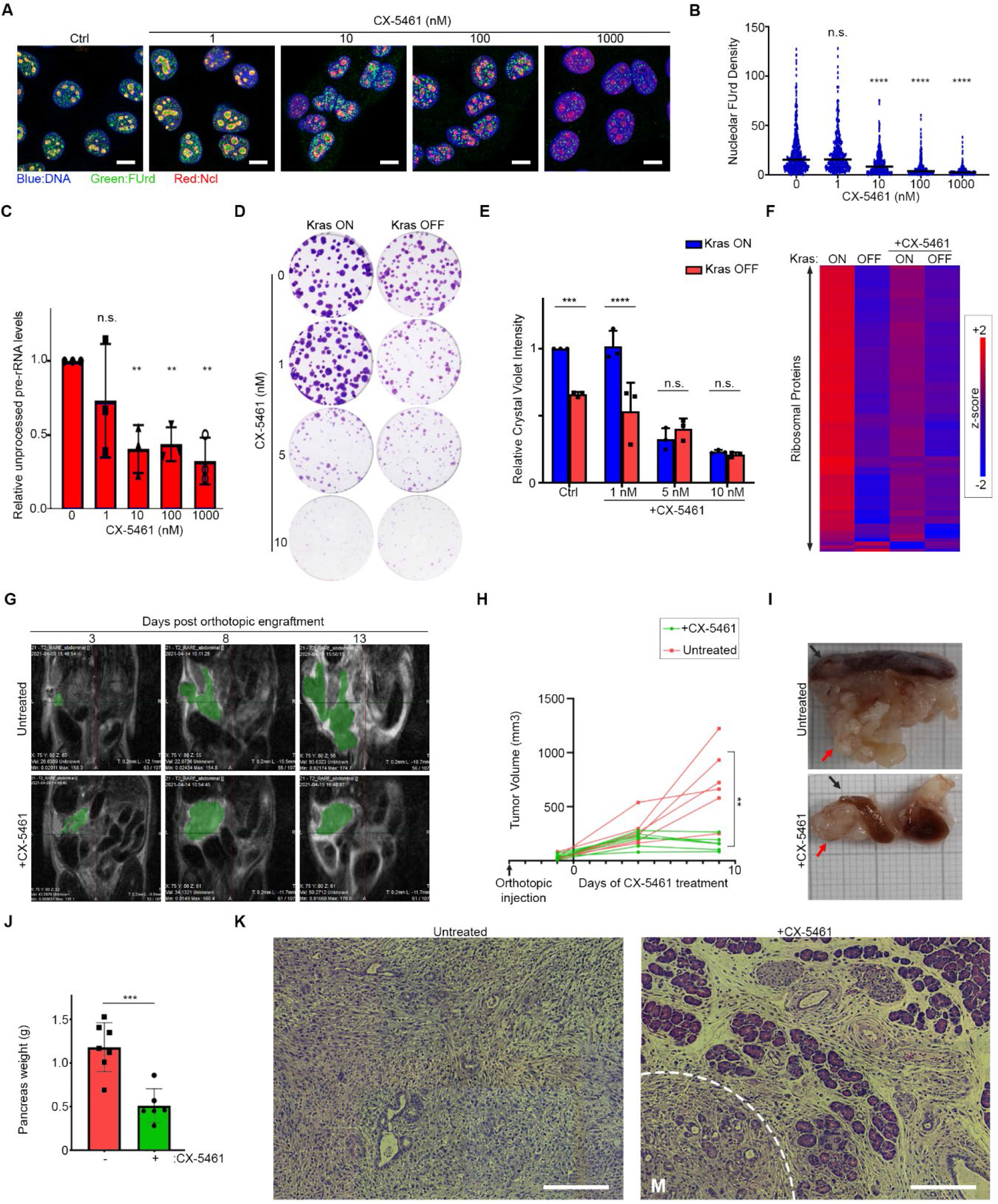
PDAC dependency on Kras^G12D^-induced ribosome biogenesis can be therapeutically targeted. **(A)** Dose response analysis of CX-5461 impact on nascent rRNA expression. IKras PDAC cells grown in presence of Dox were pre-treated with the indicated concentrations of CX-5461 for 30 mins, before pulse labeling with FUrd. Cells were then fixed and immunostained with anti-FUrd antibody to visualize nascent RNA (green), anti-Ncl antibody to reveal the Nucleolus (red), and Hoechst (blue) as the Nuclear stain, followed by confocal microscopy analysis. Scale bar = 10 μm. **(B)** Quantification of Nucleolar FUrd levels in images from (A). FUrd fluorescence densities in single nucleoli were quantified from 72-147 individual cells per condition, combined from two independent biological replicate experiments (****: P < 0.0001; n.s.: not significant). **(C)** Dose response analysis of CX-5461 impact on 5’ETS-containing pre-rRNA transcript levels. IKras PDAC cells grown in presence of Dox were treated overnight with the indicated doses of CX-5461, before RT-qPCR analysis with a specific probe against the mouse 5’ETS region. A probe against mouse Actb mRNA was used as loading control for normalization (n = 3) (**: P < 0.01; n.s.: not significant). **(D)** Dose response analysis of CX-5461 impact on Kras^G12D^-driven colony formation of iKras PDAC cells. IKras PDAC cells, seeded with or without Dox, were subjected to clonogenic assay in presence of the indicated doses of CX-5461 for 7 days. Colonies were visualized by Crystal Violet staining. **(E)** Quantification of Crystal Violet staining levels from (E) (n = 3). (****: P < 0.0001; ***: P < 0.001; n.s.: not significant). **(F)** Quantitative analysis of RP levels in vehicle or 100 nM CX-5461 treated iKras PDAC cells, in presence or absence of Kras^G12D^. IKras PDAC cells were grown for 48hrs, with or without Dox, and in presence or absence of 100 nM CX-5461. Cells were then lysed and analyzed by TMT-mediated quantitative proteomics. Z-scores of TMT intensity changes for all identified RPs across the different conditions were plotted as a heat map (red → increase; blue → decrease). **(G)** MRI imaging of orthotopic iKras tumors in untreated or CX-5461 treated mice. IKras PDAC cells were engrafted into the pancreas of Dox-fed nude mice and allowed to form tumors for 4 days. Animals were then divided into two groups, with the first group treated by daily oral administration of CX-5461 (50mg/kg) for a further 10 days, whilst the second group was left untreated for the same period. T2 scans were taken on the indicated days, post-engraftment. Green areas mark the tumors. **(H)** Quantification of tumor volumes from (G). For each condition, six animals were engrafted, treated, and analyzed by MRI imaging. (**: P < 0.01; n.s.: not significant). **(I)** Representative images of the pancreas (red arrows) from untreated and CX-5461 treated mice in (G), extracted at the end of the treatment (day 14). Spleen (black arrow), which is located adjacent to the pancreas, was also included in the images for comparison. **(J)** Quantification of pancreas weights from (I). Pancreas weight from each animal in the untreated vs. treated group was measured after extraction. (***: P < 0.001; n.s.: not significant). **(K)** H&E analysis of the extracted pancreas tissues from (I). Representative pancreatic tissue images from the untreated and CX-5461 treated mice, showing typical PDAC histology with malignant ductal structures surrounded by stroma in the untreated, but a largely normal pancreas histology with a small PDAC component (region marked M) in the treated mice. Scale bar = 200 μm.

Next, we evaluated the impact of CX-5461 on the proteome of iKras PDAC cells. Similar to Kras^G12D^ removal, nanomolar dose treatment of CX-5461 induced a strong decrease in the levels of Ribosome and rRNA processing protein categories, without affecting protein categories involved in the DNA damage response (Figure S6C & Dataset S12). Accordingly, when combined with Kras^G12D^ induction, this CX-5461 treatment could suppress the Kras^G12D^-induced accumulation of RPs (Figure 6F). Collectively, these results suggest that nanomolar doses of CX-5461 primarily inhibit the iKras PDAC cell proliferation via inhibition of Kras^G12D^-driven ribosome biogenesis, with significant DNA damage occurring only at higher doses.

Finally, we assessed whether pharmacological inhibition of ribosome biogenesis by CX-5461 could inhibit Kras^G12D^-induced tumor growth, *in vivo*. To mimic therapeutic settings, orthotopic xenograft tumors were first established in doxycycline-fed mice for 4 days, before the animals were subjected to treatment by oral administration of CX-5461 for 10 days, or left untreated for the same period as control. Magnetic Resonance Imaging (MRI) was used to monitor disease progression for the duration of the experiment. MRI imaging revealed that while tumors grew rapidly in untreated mice, tumor growth was halted in response to CX-5461 treatment (Figure 6G & 6H). End-point analysis of the pancreatic tissues also revealed a significant inhibition of tumor growth in CX-5461 treated mice, with histological analysis showing a strong reduction in the proportion of the malignant component in comparison to untreated animals (Figure 6I-K). Immunohistochemistry (IHC) analysis revealed no evidence of DNA damage in the tumors, suggesting that the *in vivo* CX-5461 dose used in our study (50 mg/kg) was comparable with the *in vitro* low dose treatments that do not induce DNA damage (Figure S6D). Together, these results reveal that pharmacological inhibition of ribosome biogenesis, the critical downstream target of the Kras^G12D^-Ncl axis, could be used as a therapeutic strategy for inhibiting PDAC growth.

## Discussion

As central modulators of post-transcriptional regulation, many RBPs have been shown to play key roles in cancer development and progression (Kang *et al*., 2020). Direct alteration of RBPs by mutation is relatively rare in cancer (Gebauer *et al*, 2021), so a key question concerns the mechanistic link between mutations in cancer driver genes and the resulting dysregulation of RBPs. In this study, we used an inducible mouse model of PDAC, combined with a whole-transcriptome quantitative RIC approach, to unbiasedly assess changes in the RBPome in response to induction of oncogenic RAS signaling. Our results reveal a drastic rewiring of the RBPome upon Kras^G12D^ induction, with many conventional RBPs showing an increase in their association with RNA, whilst non-conventional RBPs exhibit decreased association. This switch occurs downstream of ERK1/2, and is achieved not only through modulation of the expression of RBPs, but also their RNA-binding activity. In particular, we reveal a network of nuclear RBPs that include Ncl, whose RNA-binding activity increases upon Kras^G12D^ induction. Several of these RBPs undergo phosphorylation downstream of ERK1/2, and in the case of Ncl, we demonstrate that these phosphorylations act to enhance the RNA-binding activity. On the other hand, little phosphorylation changes were observed on non-conventional RBPs whose RNA-binding activity was reduced upon Kras^G12D^ induction, suggesting that other mechanisms must be at play in regulating their activity. It is tempting to speculate that other types of post-translational modifications may be involved in modulating the activity of these RBPs, but further work will be necessary to define the mechanisms of regulation, as well as the functional significance of these RBPs in the context of oncogenic RAS signaling. Nevertheless, our study demonstrates that the RNA-binding activity of many conventional and non-conventional RBPs is highly dynamic and subject to regulation by oncogenic signal transduction pathways.

Ribosome biogenesis is a highly coordinated cellular process, which involves stepwise synthesis, processing, modification, and assembly of rRNA and RPs into mature ribosomal subunits (Pelletier *et al*., 2018). Ncl is known to play a key role in several steps of ribosome biogenesis, from promotion of pre-rRNA synthesis (Cong *et al*, 2012; Roger *et al*, 2003), to mediating the first steps of pre-rRNA processing (Ginisty *et al*, 1998), as well as loading of ribosomal proteins onto pre-rRNA (Bouvet *et al*, 1998). The central RRM domains of Ncl, which mediate its binding to RNA, have been shown to be important for enhancing nascent pre-rRNA expression (Storck *et al*, 2009), suggesting that this enhancement must be at least in part achieved post-transcriptionally. In line with these findings, our results reveal that oncogenic RAS signaling activates the RNA binding activity of Ncl through phosphorylation, leading to enhancement of pre-rRNA expression, and promotion of ribosome biogenesis (Figure 7). It is not yet clear how Ncl could act to upregulate pre-rRNA expression post-transcriptionally, but structural studies suggest that Ncl may act as an RNA chaperone to mediate the proper folding of pre-rRNA (Allain *et al*, 2000). Interestingly, recent data suggests that such nascent RNA folding could be crucial for enhancing RNA polymerase-I elongation rate by inhibiting polymerase backtracking (Turowski *et al*, 2020), thus providing a possible mechanism for post-transcriptional enhancement of pre-rRNA synthesis via Ncl’s binding to pre-rRNA.

**Figure 7:**
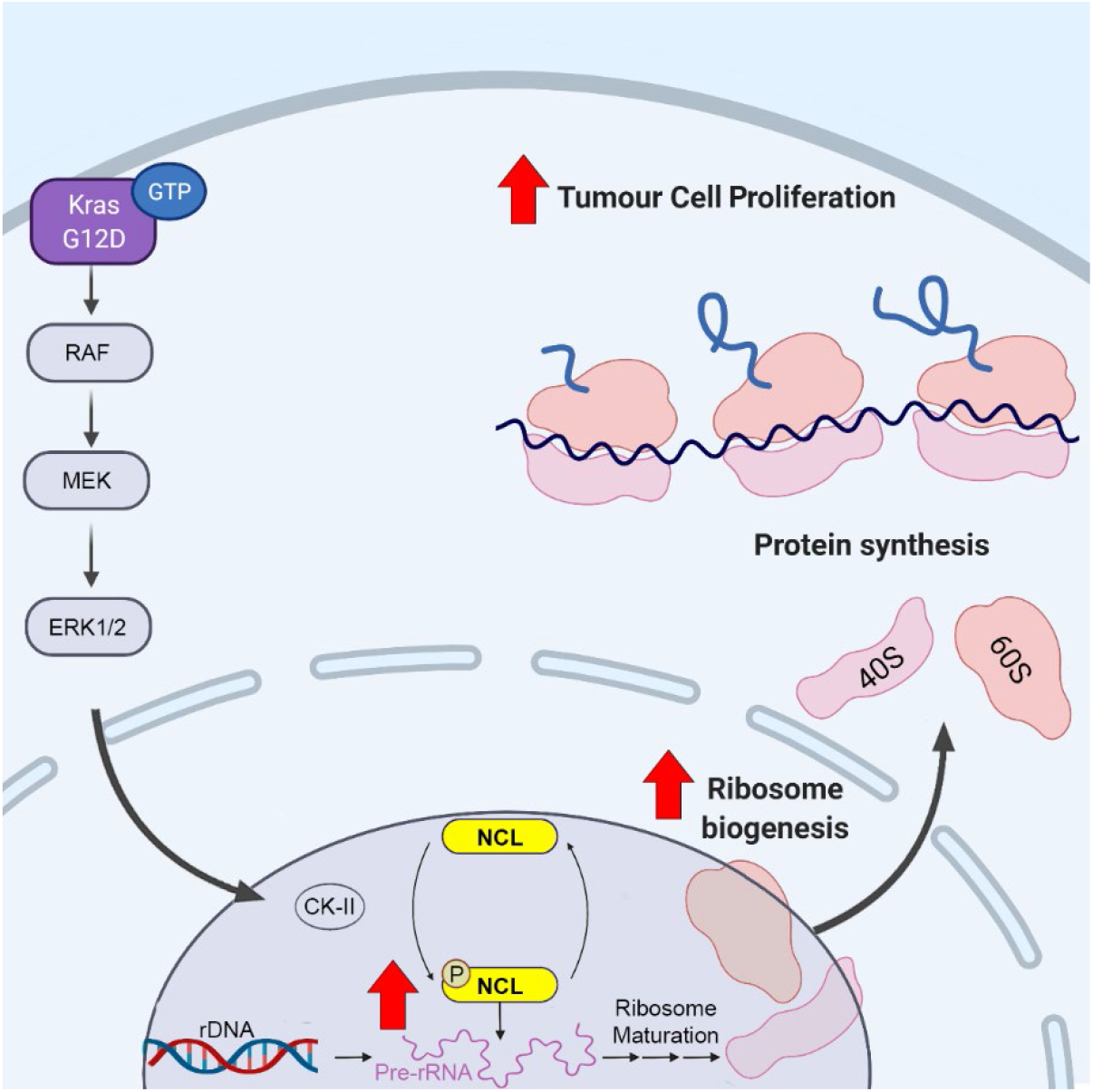
Proposed mechanism of ribosome biogenesis upregulation by oncogenic RAS signaling, via NCL. Oncogenic RAS-induced ERK activity acts to enhance CK2 activity, leading to increased phosphorylation of NCL on specific Serine residues within its N-terminal region. These phosphorylations increase the RNA-binding activity of NCL, resulting in its enhanced pre-rRNA binding, and accumulation of nascent pre-rRNA. Higher levels of pre-rRNA ultimately enhance ribosome production, leading to increased tumor cell proliferation in RAS-driven cancers.

Importantly, we revealed that while the expression of the phospho-mimicking Ncl mutant could rescue nascent pre-rRNA expression in the absence of RAS oncogene, downstream processing of pre-rRNA was blocked, indicating that the activity of other RAS-driven factors are likely to be crucial for mediating rRNA processing. Accordingly, our qRIC analysis revealed that in addition to Ncl, the RNA binding activity of several other ribosome biogenesis factors were induced upon oncogenic RAS signaling. Whether any of these factors could be playing a role in mediating other key steps of ribosome biogenesis downstream of RAS, remains to be determined. Nevertheless, our findings reveal that phosphorylation of Ncl downstream of RAS plays a key role in mediating rRNA synthesis, which is the crucial initiating step in the process of ribosome biogenesis.

Hyperactive ribosome biogenesis is a common feature of a wide variety of human cancers, playing a pivotal role in sustaining the growth and proliferation of cancer cells (Pelletier *et al*., 2018). Pre-rRNA synthesis is considered to be the rate-limiting step in the process of human ribosome biogenesis (Lam *et al*., 2007). However, mechanisms by which cancer cells upregulate pre-rRNA production are only beginning to be characterized (Arabi *et al*, 2005; Delloye-Bourgeois *et al*, 2012; Hannan *et al*, 2003; Justilien *et al*, 2017; Prakash *et al*., 2019; Zhao *et al*, 2003). Here we reveal that through phospho-regulation of Ncl, oncogenic RAS signaling enhances pre-rRNA expression and ribosome biogenesis. This upregulation is crucial for mediating PDAC cell proliferation and tumorigenesis. Accordingly, depletion of Ncl, or pharmacological inhibition of pre-rRNA synthesis by CX-5461, results in abrogation of Kras^G12D^-induced ribosome biogenesis, leading to inhibition of PDAC cell proliferation and tumor growth. In agreement with a previous report (Prakash *et al*., 2019), we confirm that when used at low nanomolar doses, the anti-proliferative effects of CX-5461 are mediated by inhibition of ribosome biogenesis, and not DNA damage induction, which can be caused at higher doses in our model. Based on these results, targeting ribosome biogenesis, either alone or in combination with other targeted therapies, appears to be a promising therapeutic avenue against RAS-driven cancers. CX-5461 has shown promise in early clinical trials (Hilton *et al*., 2018; Khot *et al*., 2019), and while our results suggest that its use could be therapeutically beneficial against PDAC, extra care must be taken in choosing the treatment dosage if DNA damage-related side effects are to be avoided. Alternatively, more specific inhibitors of rRNA synthesis may become available in the near future, whose therapeutic effectiveness against RAS-driven cancers warrants further investigation. In addition, direct inhibition of nucleolar Ncl, or its phospho-regulation downstream of RAS, could be another attractive strategy for targeting ribosome biogenesis in RAS-driven tumors.

## Experimental procedures

Data availability information, full list of reagents, and detailed experimental procedures are available in the Supplementary Information.

## Author contributions

<u>F.K.M.</u> conceived the study, supervised the work, and wrote the manuscript. <u>M.Dodel.</u> performed the mouse experiments, and the qRIC analyses. <u>F.C.</u> performed the iCLIP experiment. <u>R.F.</u> and <u>J.U.</u> analyzed the iCLIP data. <u>M.Dermit</u> carried out the phospho-proteomics analysis. <u>W.F.</u> assisted with the CK2 activity assessments. <u>M.S.A</u> performed all other experiments and analyzed the data.

## Acknowledgements

This work was funded and supported by a Medical Research Council Career Development Award (MR/P009417/1) and a Barts Charity grant (MGU0346) to <u>F.K.M.</u>, a Barry Reed PhD studentship to <u>M.S.A.</u>, and a Cancer Research UK Centre Grant to Barts Cancer Institute (C355/A25137). We would like to acknowledge the Pre-clinical Imaging facility, the Animal Technician Services (ATS), and the Biological Services Unit (BSU) at Barts Cancer Institute, for their substantial support with the animal work. We would also like to acknowledge the Microscopy, Histopathology, and Mass spectrometry core facilities for their support with imaging and proteomics experiments, and the Barts and the London Genome Centre for the next-generation sequencings. Finally, we would like to thank Dr. Gunnel Hallden for her valuable input on animal experiments, and Dr. Christopher Tape for sharing of reagents and advice on the project.

## References

Abdelmohsen K, Gorospe M (2012) RNA-binding protein nucleolin in disease. RNA Biol 9: 799–808

Abdelmohsen K, Tominaga K, Lee EK, Srikantan S, Kang MJ, Kim MM, Selimyan R, Martindale JL, Yang X, Carrier F et al (2011) Enhanced translation by Nucleolin via G-rich elements in coding and non-coding regions of target mRNAs. Nucleic acids research 39: 8513–8530

Allain FH, Bouvet P, Dieckmann T, Feigon J (2000) Molecular basis of sequence-specific recognition of pre-ribosomal RNA by nucleolin. The EMBO journal 19: 6870–6881

Angelov D, Bondarenko VA, Almagro S, Menoni H, Mongelard F, Hans F, Mietton F, Studitsky VM, Hamiche A, Dimitrov S et al (2006) Nucleolin is a histone chaperone with FACT-like activity and assists remodeling of nucleosomes. The EMBO journal 25: 1669–1679

Arabi A, Wu S, Ridderstrale K, Bierhoff H, Shiue C, Fatyol K, Fahlen S, Hydbring P, Soderberg O, Grummt I et al (2005) c-Myc associates with ribosomal DNA and activates RNA polymerase I transcription. Nature cell biology 7: 303–310

Baltz AG, Munschauer M, Schwanhausser B, Vasile A, Murakawa Y, Schueler M, Youngs N, Penfold-Brown D, Drew K, Milek M et al (2012) The mRNA-bound proteome and its global occupancy profile on protein-coding transcripts. Molecular cell 46: 674–690

Barbieri I, Tzelepis K, Pandolfini L, Shi J, Millan-Zambrano G, Robson SC, Aspris D, Migliori V, Bannister AJ, Han N et al (2017) Promoter-bound METTL3 maintains myeloid leukaemia by m(6)A-dependent translation control. Nature 552: 126–131

Bouvet P, Diaz JJ, Kindbeiter K, Madjar JJ, Amalric F (1998) Nucleolin interacts with several ribosomal proteins through its RGG domain. J Biol Chem 273: 19025–19029

Bruno PM, Lu M, Dennis KA, Inam H, Moore CJ, Sheehe J, Elledge SJ, Hemann MT, Pritchard JR (2020) The primary mechanism of cytotoxicity of the chemotherapeutic agent CX-5461 is topoisomerase II poisoning. Proceedings of the National Academy of Sciences of the United States of America 117: 4053–4060

Bywater MJ, Poortinga G, Sanij E, Hein N, Peck A, Cullinane C, Wall M, Cluse L, Drygin D, Anderes K et al (2012) Inhibition of RNA polymerase I as a therapeutic strategy to promote cancer-specific activation of p53. Cancer Cell 22: 51–65

Caizergues-Ferrer M, Belenguer P, Lapeyre B, Amalric F, Wallace MO, Olson MO (1987) Phosphorylation of nucleolin by a nucleolar type NII protein kinase. Biochemistry 26: 7876–7883

Castello A, Fischer B, Eichelbaum K, Horos R, Beckmann BM, Strein C, Davey NE, Humphreys DT, Preiss T, Steinmetz LM et al (2012) Insights into RNA biology from an atlas of mammalian mRNA-binding proteins. Cell 149: 1393–1406

Chamousset D, De Wever V, Moorhead GB, Chen Y, Boisvert FM, Lamond AI, Trinkle-Mulcahy L (2010) RRP1B targets PP1 to mammalian cell nucleoli and is associated with Pre-60S ribosomal subunits. Mol Biol Cell 21: 4212–4226

Cong R, Das S, Ugrinova I, Kumar S, Mongelard F, Wong J, Bouvet P (2012) Interaction of nucleolin with ribosomal RNA genes and its role in RNA polymerase I transcription. Nucleic acids research 40: 9441–9454

de Beus E, Brockenbrough JS, Hong B, Aris JP (1994) Yeast NOP2 encodes an essential nucleolar protein with homology to a human proliferation marker. The Journal of cell biology 127: 1799–1813

Delloye-Bourgeois C, Goldschneider D, Paradisi A, Therizols G, Belin S, Hacot S, Rosa-Calatrava M, Scoazec JY, Diaz JJ, Bernet A et al (2012) Nucleolar localization of a netrin-1 isoform enhances tumor cell proliferation. Sci Signal 5: ra57

Dermit M, Dodel M, Lee FCY, Azman MS, Schwenzer H, Jones JL, Blagden SP, Ule J, Mardakheh FK (2020) Subcellular mRNA Localization Regulates Ribosome Biogenesis in Migrating Cells. Developmental cell 55: 298–313 e210

Drygin D, Lin A, Bliesath J, Ho CB, O’Brien SE, Proffitt C, Omori M, Haddach M, Schwaebe MK, Siddiqui-Jain A et al (2011) Targeting RNA polymerase I with an oral small molecule CX-5461 inhibits ribosomal RNA synthesis and solid tumor growth. Cancer research 71: 1418–1430

Erard MS, Belenguer P, Caizergues-Ferrer M, Pantaloni A, Amalric F (1988) A major nucleolar protein, nucleolin, induces chromatin decondensation by binding to histone H1. Eur J Biochem 175: 525–530

Fish L, Khoroshkin M, Navickas A, Garcia K, Culbertson B, Hanisch B, Zhang S, Nguyen HCB, Soto LM, Dermit M et al (2021) A prometastatic splicing program regulated by SNRPA1 interactions with structured RNA elements. Science 372

Franceschini A, Szklarczyk D, Frankild S, Kuhn M, Simonovic M, Roth A, Lin J, Minguez P, Bork P, von Mering C et al (2013) STRING v9.1: protein-protein interaction networks, with increased coverage and integration. Nucleic acids research 41: D808–815

Garcia-Moreno M, Noerenberg M, Ni S, Jarvelin AI, Gonzalez-Almela E, Lenz CE, Bach-Pages M, Cox V, Avolio R, Davis T et al (2019) System-wide Profiling of RNA-Binding Proteins Uncovers Key Regulators of Virus Infection. Molecular cell 74: 196–211 e111

Gebauer F, Schwarzl T, Valcarcel J, Hentze MW (2021) RNA-binding proteins in human genetic disease. Nature reviews Genetics 22: 185–198

Gerstberger S, Hafner M, Tuschl T (2014) A census of human RNA-binding proteins. Nature reviews Genetics 15: 829–845

Ginisty H, Amalric F, Bouvet P (1998) Nucleolin functions in the first step of ribosomal RNA processing. The EMBO journal 17: 1476–1486

Hannan KM, Brandenburger Y, Jenkins A, Sharkey K, Cavanaugh A, Rothblum L, Moss T, Poortinga G, McArthur GA, Pearson RB et al (2003) mTOR-dependent regulation of ribosomal gene transcription requires S6K1 and is mediated by phosphorylation of the carboxy-terminal activation domain of the nucleolar transcription factor UBF. Molecular and cellular biology 23: 8862–8877

Hentze MW, Castello A, Schwarzl T, Preiss T (2018) A brave new world of RNA-binding proteins. Nature reviews Molecular cell biology 19: 327–341

Hilton J, Cescon DW, Bedard P, Ritter H, Tu D, Soong J, Gelmon K, Aparicio S, Seymour L (2018) 44O - CCTG IND.231: A phase 1 trial evaluating CX-5461 in patients with advanced solid tumors. Annals of Oncology 29: iii8

Hong S, Freeberg MA, Han T, Kamath A, Yao Y, Fukuda T, Suzuki T, Kim JK, Inoki K (2017) LARP1 functions as a molecular switch for mTORC1-mediated translation of an essential class of mRNAs. Elife 6

Ji H, Wang J, Nika H, Hawke D, Keezer S, Ge Q, Fang B, Fang X, Fang D, Litchfield DW et al (2009) EGF-induced ERK activation promotes CK2-mediated disassociation of alpha-Catenin from beta-Catenin and transactivation of beta-Catenin. Molecular cell 36: 547–559

Jumper J, Evans R, Pritzel A, Green T, Figurnov M, Ronneberger O, Tunyasuvunakool K, Bates R, Zidek A, Potapenko A et al (2021) Highly accurate protein structure prediction with AlphaFold. Nature 596: 583–589

Justilien V, Ali SA, Jamieson L, Yin N, Cox AD, Der CJ, Murray NR, Fields AP (2017) Ect2-Dependent rRNA Synthesis Is Required for KRAS-TRP53-Driven Lung Adenocarcinoma. Cancer Cell 31: 256–269

Kang D, Lee Y, Lee JS (2020) RNA-Binding Proteins in Cancer: Functional and Therapeutic Perspectives. Cancers (Basel) 12

Khot A, Brajanovski N, Cameron DP, Hein N, Maclachlan KH, Sanij E, Lim J, Soong J, Link E, Blombery P et al (2019) First-in-Human RNA Polymerase I Transcription Inhibitor CX-5461 in Patients with Advanced Hematologic Cancers: Results of a Phase I Dose-Escalation Study. Cancer Discov 9: 1036–1049

Lam YW, Lamond AI, Mann M, Andersen JS (2007) Analysis of nucleolar protein dynamics reveals the nuclear degradation of ribosomal proteins. Current biology: CB 17: 749–760

Li D, Dobrowolska G, Krebs EG (1996) The physical association of casein kinase 2 with nucleolin. J Biol Chem 271: 15662–15668

Linding R, Jensen LJ, Pasculescu A, Olhovsky M, Colwill K, Bork P, Yaffe MB, Pawson T (2008) NetworKIN: a resource for exploring cellular phosphorylation networks. Nucleic acids research 36: D695–699

Liu Y, Mattila J, Ventela S, Yadav L, Zhang W, Lamichane N, Sundstrom J, Kauko O, Grenman R, Varjosalo M et al (2017) PWP1 Mediates Nutrient-Dependent Growth Control through Nucleolar Regulation of Ribosomal Gene Expression. Developmental cell 43: 240–252 e245

Losfeld ME, Khoury DE, Mariot P, Carpentier M, Krust B, Briand JP, Mazurier J, Hovanessian AG, Legrand D (2009) The cell surface expressed nucleolin is a glycoprotein that triggers calcium entry into mammalian cells. Exp Cell Res 315: 357–369

Malumbres M, Barbacid M (2003) RAS oncogenes: the first 30 years. Nat Rev Cancer 3: 459–465

Mars JC, Tremblay MG, Valere M, Sibai DS, Sabourin-Felix M, Lessard F, Moss T (2020) The chemotherapeutic agent CX-5461 irreversibly blocks RNA polymerase I initiation and promoter release to cause nucleolar disruption, DNA damage and cell inviability. NAR Cancer 2: zcaa032

Matter N, Herrlich P, Konig H (2002) Signal-dependent regulation of splicing via phosphorylation of Sam68. Nature 420: 691–695

McAlister GC, Huttlin EL, Haas W, Ting L, Jedrychowski MP, Rogers JC, Kuhn K, Pike I, Grothe RA, Blethrow JD et al (2012) Increasing the multiplexing capacity of TMTs using reporter ion isotopologues with isobaric masses. Analytical chemistry 84: 7469–7478

Moudry P, Chroma K, Bursac S, Volarevic S, Bartek J (2021) RNA-interference screen for p53 regulators unveils a role of WDR75 in ribosome biogenesis. Cell Death Differ

Negi SS, Brown P (2015) rRNA synthesis inhibitor, CX-5461, activates ATM/ATR pathway in acute lymphoblastic leukemia, arrests cells in G2 phase and induces apoptosis. Oncotarget 6: 18094–18104

Nusinow DP, Szpyt J, Ghandi M, Rose CM, McDonald ER, 3rd, Kalocsay M, Jane-Valbuena J, Gelfand E, Schweppe DK, Jedrychowski M et al (2020) Quantitative Proteomics of the Cancer Cell Line Encyclopedia. Cell 180: 387–402 e316

Ong SE, Mann M (2006) A practical recipe for stable isotope labeling by amino acids in cell culture (SILAC). Nat Protoc 1: 2650–2660

Pan M, Wright WC, Chapple RH, Zubair A, Sandhu M, Batchelder JE, Huddle BC, Low J, Blankenship KB, Wang Y et al (2021) The chemotherapeutic CX-5461 primarily targets TOP2B and exhibits selective activity in high-risk neuroblastoma. Nat Commun 12: 6468

Pelletier J, Thomas G, Volarevic S (2018) Ribosome biogenesis in cancer: new players and therapeutic avenues. Nat Rev Cancer 18: 51–63

Percipalle P, Louvet E (2012) In vivo run-on assays to monitor nascent precursor RNA transcripts. Methods in molecular biology 809: 519–533

Pereira B, Billaud M, Almeida R (2017) RNA-Binding Proteins in Cancer: Old Players and New Actors. Trends Cancer 3: 506–528

Pichiorri F, Palmieri D, De Luca L, Consiglio J, You J, Rocci A, Talabere T, Piovan C, Lagana A, Cascione L et al (2013) In vivo NCL targeting affects breast cancer aggressiveness through miRNA regulation. J Exp Med 210: 951–968

Pickering BF, Yu D, Van Dyke MW (2011) Nucleolin protein interacts with microprocessor complex to affect biogenesis of microRNAs 15a and 16. J Biol Chem 286: 44095–44103

Pineiro D, Stoneley M, Ramakrishna M, Alexandrova J, Dezi V, Juke-Jones R, Lilley KS, Cain K, Willis AE (2018) Identification of the RNA polymerase I-RNA interactome. Nucleic acids research 46: 11002–11013

Prakash V, Carson BB, Feenstra JM, Dass RA, Sekyrova P, Hoshino A, Petersen J, Guo Y, Parks MM, Kurylo CM et al (2019) Ribosome biogenesis during cell cycle arrest fuels EMT in development and disease. Nat Commun 10: 2110

Prior IA, Hood FE, Hartley JL (2020) The Frequency of Ras Mutations in Cancer. Cancer research 80: 2969–2974

Pylayeva-Gupta Y, Grabocka E, Bar-Sagi D (2011) RAS oncogenes: weaving a tumorigenic web. Nat Rev Cancer 11:761–774

Queiroz RML, Smith T, Villanueva E, Marti-Solano M, Monti M, Pizzinga M, Mirea DM, Ramakrishna M, Harvey RF, Dezi V et al (2019) Comprehensive identification of RNA-protein interactions in any organism using orthogonal organic phase separation (OOPS). Nat Biotechnol 37: 169–178

Quin J, Chan KT, Devlin JR, Cameron DP, Diesch J, Cullinane C, Ahern J, Khot A, Hein N, George AJ et al (2016) Inhibition of RNA polymerase I transcription initiation by CX-5461 activates non-canonical ATM/ATR signaling. Oncotarget 7: 49800–49818

Reyes-Reyes EM, Akiyama SK (2008) Cell-surface nucleolin is a signal transducing P-selectin binding protein for human colon carcinoma cells. Exp Cell Res 314: 2212–2223

Roger B, Moisand A, Amalric F, Bouvet P (2003) Nucleolin provides a link between RNA polymerase I transcription and pre-ribosome assembly. Chromosoma 111: 399–407

Sanij E, Hannan KM, Xuan J, Yan S, Ahern JE, Trigos AS, Brajanovski N, Son J, Chan KT, Kondrashova O et al (2020) CX-5461 activates the DNA damage response and demonstrates therapeutic efficacy in high-grade serous ovarian cancer. Nat Commun 11: 2641

Sengupta TK, Bandyopadhyay S, Fernandes DJ, Spicer EK (2004) Identification of nucleolin as an AU-rich element binding protein involved in bcl-2 mRNA stabilization. J Biol Chem 279: 10855–10863

Storck S, Thiry M, Bouvet P (2009) Conditional knockout of nucleolin in DT40 cells reveals the functional redundancy of its RNA-binding domains. Biol Cell 101: 153–167

Sysoev VO, Fischer B, Frese CK, Gupta I, Krijgsveld J, Hentze MW, Castello A, Ephrussi A (2016) Global changes of the RNA-bound proteome during the maternal-to-zygotic transition in Drosophila. Nat Commun 7: 12128

Tajrishi MM, Tuteja R, Tuteja N (2011) Nucleolin: The most abundant multifunctional phosphoprotein of nucleolus. Commun Integr Biol 4: 267–275

Takagi M, Absalon MJ, McLure KG, Kastan MB (2005) Regulation of p53 translation and induction after DNA damage by ribosomal protein L26 and nucleolin. Cell 123: 49–63

Tape CJ, Ling S, Dimitriadi M, McMahon KM, Worboys JD, Leong HS, Norrie IC, Miller CJ, Poulogiannis G, Lauffenburger DA et al (2016) Oncogenic KRAS Regulates Tumor Cell Signaling via Stromal Reciprocation. Cell 165: 910–920

Trendel J, Schwarzl T, Horos R, Prakash A, Bateman A, Hentze MW, Krijgsveld J (2019) The Human RNA-Binding Proteome and Its Dynamics during Translational Arrest. Cell 176: 391–403 e319

Tripathi V, Ellis JD, Shen Z, Song DY, Pan Q, Watt AT, Freier SM, Bennett CF, Sharma A, Bubulya PA et al (2010) The nuclear-retained noncoding RNA MALAT1 regulates alternative splicing by modulating SR splicing factor phosphorylation. Molecular cell 39: 925–938

Truitt ML, Conn CS, Shi Z, Pang X, Tokuyasu T, Coady AM, Seo Y, Barna M, Ruggero D (2015) Differential Requirements for eIF4E Dose in Normal Development and Cancer. Cell 162: 59–71

Turowski TW, Petfalski E, Goddard BD, French SL, Helwak A, Tollervey D (2020) Nascent Transcript Folding Plays a Major Role in Determining RNA Polymerase Elongation Rates. Molecular cell 79: 488–503 e411

Turowski TW, Tollervey D (2015) Cotranscriptional events in eukaryotic ribosome synthesis. Wiley interdisciplinary reviews RNA 6: 129–139

Ugrinova I, Petrova M, Chalabi-Dchar M, Bouvet P (2018) Multifaceted Nucleolin Protein and Its Molecular Partners in Oncogenesis. Adv Protein Chem Struct Biol 111: 133–164

Waters AM, Der CJ (2018) KRAS: The Critical Driver and Therapeutic Target for Pancreatic Cancer. Cold Spring Harb Perspect Med 8

Wright CJ, McCormack PL (2013) Trametinib: first global approval. Drugs 73: 1245–1254

Xu H, Di Antonio M, McKinney S, Mathew V, Ho B, O’Neil NJ, Santos ND, Silvester J, Wei V, Garcia J et al (2017) CX-5461 is a DNA G-quadruplex stabilizer with selective lethality in BRCA1/2 deficient tumours. Nat Commun 8: 14432

Ying H, Kimmelman AC, Lyssiotis CA, Hua S, Chu GC, Fletcher-Sananikone E, Locasale JW, Son J, Zhang H, Coloff JL et al (2012) Oncogenic Kras maintains pancreatic tumors through regulation of anabolic glucose metabolism. Cell 149: 656–670

Yoon S, Seger R (2006) The extracellular signal-regulated kinase: multiple substrates regulate diverse cellular functions. Growth Factors 24: 21–44

Yu J, Navickas A, Asgharian H, Culbertson B, Fish L, Garcia K, Olegario JP, Dermit M, Dodel M, Hanisch B et al (2020) RBMS1 Suppresses Colon Cancer Metastasis through Targeted Stabilization of Its mRNA Regulon. Cancer Discov 10: 1410–1423

Zhang Y, Bhatia D, Xia H, Castranova V, Shi X, Chen F (2006) Nucleolin links to arsenic-induced stabilization of GADD45alpha mRNA. Nucleic acids research 34: 485–495

Zhao J, Yuan X, Frodin M, Grummt I (2003) ERK-dependent phosphorylation of the transcription initiation factor TIF-IA is required for RNA polymerase I transcription and cell growth. Molecular cell 11: 405–413

