## Supplementary Information for "An RNA-binding switch drives ribosome biogenesis and tumorigenesis downstream of RAS oncogene"

Supplementary information for this manuscript consists of Supplementary Experimental Procedures, Supplementary Figures (S1 to S6), Supplementary References, and a supplemental Excel file containing Supplementary Datasets (S1 to S12).

### Supplementary experimental procedures

#### Plasmids and Reagents

WT and mutant Myc-Ncl expression plasmids were generated by Gateway cloning of custom synthesized murine Ncl donor vectors (GeneArt) into the Myc-pRK5-DEST vector. Detailed list of all other reagents used in this study can be found in the Key Resources Table below.

#### Cell-culture and transfections

Mouse iKras PDAC cells were grown in DMEM supplemented with 10% FBS, 1% Penicillin/Streptomycin, and doxycycline (1µg/ml). To remove $Kras^{G12D}$ expression, the same media without doxycycline was used. Cells were grown in humidified incubator at 37°C with 5% CO2, and routinely checked to be mycoplasma-free by MycoAlert Plus mycoplasma detection kit. Collagen-I matrix was prepared as described previously (Dermit et al., 2020), and used for assessment of cell growth in 3D. For siRNA-mediated depletions, 10,000 cells/cm$^2$ were seeded on standard TC-treated polystyrene plates and transfected the next day using Lipofectamine RNAiMAX (Thermo), according to manufacturer's instructions. A final siRNA concentration of 20 nM was used per condition. For efficient Ncl depletion, cells were double or triple transfected at 48 hr intervals. For DNA transfections, 25,000 cells/cm$^2$ were seeded on standard TC-treated polystyrene plates and transfected the next day using Lipofectamine 2000 (Thermo), according to manufacturer's instructions. A final DNA amount of 250 ng/cm$^2$ was used per condition, and cells were re-seeded the next day for downstream analyses.

#### Immunofluorescence (IF)

IF was carried out in 18 Well Flat µ-Slides from iBidi. 2,000 cells were seeded in each well and grown for 48 hrs in presence or absence of doxycycline (1µg/ml), before the indicated treatments. Cells were washed with PBS and fixed in fixation buffer (4% Formaldehyde in PBS) for 15 min. The fixed cells were then permeabilized with permeabilization buffer (0.5% Triton-X100 in PBS) for 10 minutes before 3x washes with PBS. Cells were then incubated with blocking buffer (4%BSA in PBS) for 30 minutes, before incubation with the indicated primary antibodies (diluted in blocking buffer) for 1 hr at room temperature (RT). This was followed by 3X PBS washes and incubation with fluorophore-conjugated secondary antibodies (diluted in blocking buffer) for another hour at RT in the dark. Slides were then washed again 3X with PBS and imaged on a Zeiss LSM 880 confocal microscope using a 63X oil immersion lens. FUrd pulse labeling was done according to (Percipalle and Louvet, 2012), with some modifications. Briefly, cells were pulsed for 30 minutes with 2mM FUrd before fixation. RNAse-free reagents were used for preparation of fixation, permeabilization, and blocking buffers, and the blocking buffer was supplemented with SUPERase-In RNase Inhibitor (Thermo) at 1:500 dilution, to inhibit RNA degradation. To visualize FUrd incorporation, a monoclonal antibody against BrdU, which also detects FUrd (B2531, Merck) was used.

**Mouse orthotopic xenograft studies**

All xenograft experiments were performed under Home Office UK Project License (PP9448177), protocol 3, after internal review board approval. For orthotopic establishment of PDAC tumors, $5 \times 10^5$ iKras PDAC cells, suspended in 20 µl of 50% Matrigel (BD Biosciences)/Hanks buffered saline solution, were injected into the pancreas of NCr nude mice. Anaesthetic machine with Isoflurane for induction and duration of the procedure was used during the surgery. As analgesic, animals were given Vetergesic (buprenophine) sub cutaneous into scruff before the surgery, later in the day, and the next morning. All animals were observed and examined for any abnormalities, and body weight was measured daily. To maintain $Kras^{G12D}$ expression, animals were fed with water supplemented with 2g/L of doxycycline and 20g/L of Sucrose. CX-5461, resuspended in 50 mM sodium phosphate solution (pH 4.5) at 10 mg/ml, was administered via daily oral gavage, at a final dose of 50 mg/kg. In vivo tumor imaging was performed using a fast 3 min T2 weighted MRI scan on a Bruker ICON 1T MRI system instrument, at the indicated time-points. An anesthetic rig was used to immobilize the mice by administering isoflurane into an induction box and keeping them unconscious on the MRI bed for the duration of the imaging.

**Tissue staining**

All tissue stainings were performed by the BCI Histophatology core facility. Haematoxylin & Eosin (H&E) staining of formalin fixed paraffin embedded tissue sections was performed using the Leica Autostainer XL (V2.01). For Immunohistochemistry (IHC), a Ventana Discovery XT instrument was used for the deparaffinization, Heat Induced Epitope Retrieval, and blocking steps. Ph2AX antibody was initially tested on Cisplatin-treated and untreated iKras PDAC cells embedded in agarose, and an optimal dilution of 1:400 was determined for detection of DNA damages cells by IHC staining. Primary and secondary OmniMap HRP-conjugated antibodies were used in 100 µl volumes, for 60 and 16 minutes, respectively. Slides were then stained by applying one drop of DAB CM and One Drop $H_2O_2$ CM and incubating for 8 minutes, then applying one drop of Copper CM and incubating for 5 minutes. Slides were then counterstained with Hematoxylin, and post counterstaining with Bluing Reagent was performed, before washing with warm water with detergent, and dehydrating in graded ethanol and xylene. Slides were then covered by glass coverslips, attached with permanent mounting media.

**Image analysis**

All images were processed and analyzed using ImageJ. For analysis of nucleolar FUrd incorporation, a mask of the nucleoli was first generated using the fibrillarin or Ncl channels as nucleoli markers. For this purpose, the *Despeckle* function followed by *Smoothing* of the edges was first performed on the Fibrillarin or Ncl channels. Channel images were then converted to *Binary*, and the *Find Edges* function was used to create the nucleolar mask. Each binary nucleolus was subsequently selected as a *region of interest (ROI),* and used for quantification of the Integrated Density from the FUrd channel. For DNA damage analysis, Hoechst staining was used to generate a binary mask of the nucleus. For this purpose, the Hoechst channel image was first converted to binary, and the Find Edges function was subsequently performed to create a nuclear mask. Next, each nucleus was selected as an ROI, and the Integrated Density of the pH2AX channel was then quantified within each nucleus.

**Western Blotting**

Samples were lysed in 2% SDS, 100mM Tris/HCl pH 7.5 and sonicated with a sonicator bath (Bioruptor Pico - Rm 343) for 15 cycles. Sample concentration was adjusted with a Pierce BCA Protein Assay Kit (Thermo) before addition of NuPAGE LDS Sample Buffer (Thermo) with reducing agent and boiling at 95°C for 10 minutes. After separation on a NuPage 4%–12% Bis/Tris protein gel (Thermo), proteins were transferred to an Immobilon-P membrane (Millipore) using a standard wet transfer device. Primary antibodies were diluted in 5%BSA, PBS and incubated on the membranes at 4°C overnight followed by incubation with anti-mouse or rabbit HRP-conjugated secondary antibodies at RT for one hour. Membranes were then probed with Pierce ECL Plus HRP-detection reagent followed by imaging on an Amersham Imager 600. Western blots were quantified using ImageJ.

**Colony formation and 3D viability assays**

Colony formation assay was performed by seeding cells at 500 cells per well on a 6 well plate, with or without doxycycline (1µg/ml) for 5-7 days, with the media being replenished every 3 days. Cells were subsequently fixed with 4% formaldehyde on ice for 30 mins in the dark, followed by staining with 0.5% crystal violet staining solution (0.5% w/v, 20% MeOH) for 10 mins at RT. Plates were then imaged on an Amersham Imager 600, and ImageJ was used to quantify the crystal violet staining densities. For this purpose, images were first converted to 8-bit grey scale, recalibrated using the *Uncalibrated OD* function, and Integrated Density of then measured and quantified for each well. For 3D viability assay, cells were seeded at 20,000 cells per well of a 24 well plate, with or without doxycycline (1µg/ml) for 48 hrs. 3D Cell-Titer Glo reagent (Promega), diluted 1:4 in PBS, was then used to quantify cell viability in each well, according to the manufacturer's instructions. Luminescence was measured on a BMG Plate-reader and analyzed using GraphPad PRISM.

**RT-qPCR**

Total RNA was isolated using TRIzol reagent (Fisher), as per manufacturer's instructions. RT-qPCR was performed using Brilliant II SYBR® Green one-step kit (Agilent) on an ABI 7500 Real-Time PCR system (Applied Biosystems). The 2-ΔΔCT method was used for relative quantification of RNA expression levels, according to (Rao et al., 2013). β-actin (Actb) mRNA was also quantified and used as an internal control for normalizations. All primers used for RT-qPCR analyses are listed in the Key Resources Table.

**iCLIP**

The iCLIP method was performed as previously described in (Lee et al., 2021), with some modifications. Briefly, triplicates of mock transfected or WT Myc-Ncl, S4A Myc-Ncl, S4D Myc-Ncl transfected iKras PDAC cells were seeded onto 10 cm dishes (2 million per dish) and allowed to grow for 48 hrs in the presence of doxycycline (1µg/ml), before being irradiated once on ice with 150 mJ/cm$^2$ UV light (254 nm) in PBS, using a Hoefer Scientific UV Crosslinker. Cells were then lysed in lysis buffer (50mM Tris-HCl pH 7.4, 100mM NaCl, 1% Igepal CA-630, 0.1% SDS, 0.5% sodium deoxycholate, supplemented with protease inhibitors), cleared, and diluted to a protein concentration of 1mg/ml. RNA was then digested with 0.2 U/ml of RNase I. Myc-tagged Ncl was then immunoprecipitated with 4µg anti-Myc-tag antibody (Cell Signaling), pre-conjugated to protein G Dynabeads (Thermo). The RNAs were labelled at the 3' end using an adapter (/5Phos/AG ATC GGA AGA GCG GTT CAG AAA AAA AAA AAA/iAzideN/AA AAA AAA AAA A/3Bio/) conjugated to an infrared dye to allow the visualization of the complexes on a gel. After the SDS-PAGE and the transfer onto nitrocellulose membrane, the region corresponding to 140–200 kDa protein-RNA crosslinked complexes was excised to isolate the associated RNAs. Isolated RNAs were reverse transcribed using primers containing experimental barcodes unique to each sample (see Key Resources Table for the primer sequences). The cDNAs were then PCR amplified, gel extracted, and equal amounts of each amplified library was then combined together into a single mixed pool, before sequencing on an Illumina NextSeq 500, producing 150-nt single-end reads. For iCLIP data analysis, the reads were trimmed and demultiplexed using Ultraplex and aligned using STAR (Dobin et al., 2013; Wilkins et al., 2021). Reads were first mapped to a genome containing the ribosomal DNA repeat and other short ncRNA sequences from GENCODE vM22, before being mapped to the mm10 genome using GENCODE vM22 annotation. PCR duplicates were collapsed using UMI-tools (Smith et al., 2017). The crosslink position was defined as the nucleotide upstream of the 5' end of the read. For analysis of crosslinking to rRNA, crosslinking signal was normalized to the total number of rRNA crosslinks per sample. Proportional crosslink density was defined as the normalized crosslinks per nucleotide for each rRNA subtype.

**Orthogonal Organic Phase Separation (OOPS)**

Cells were SILAC labeled by being passaged for at least six doublings in Lysine and Arginine free DMEM, supplemented with 10% dialyzed FBS, 1% P/S, 600mg/L Proline, in the presence of 100mg/L of either light Arginine and Lysine (for ‘‘light’’ media), or heavy Arginine [U-13C6, U-15N4] and Lysine [U-13C6, U-15N2] (for ‘‘heavy” media). Experiments were always performed in duplicates, with reciprocal SILAC labelling. OOPS was carried out as described in (Queiroz et al., 2019), with some modifications. Briefly, $10^6$ heavy or light labelled iKras PDAC cells were seeded onto 10 cm dishes and grown for 48 hrs without doxycycline, before addition of doxycycline (1µg/ml) to one label, whilst leaving the other label unchanged. Cells were incubated for another 24 hrs before being washed with ice-cold PBS and irradiated on ice with 400 mJ/cm$^2$ of UV-C (254 nm), using a Hoefer Scientific UV Crosslinker. Cells were then lysed by direct addition of TRIzol reagent (Thermo) to each dish (1ml). After scraping the cells in TRIzol, the lysate from heavy and light SILAC labels were combined and homogenized through pipetting before incubating at RT for 5 min in order to dissociate non-crosslinked proteins. 200 µl of Chloroform per 1 ml of TRIZol was then added to the mix, followed by a second homogenization through vortexing. Phase separation was achieved by centrifugation at 12000g for 15 min at 4°C. After removing the organic and the aqueous phases, the interface was further purified for 3 additional times by re-dissolving in TRIZol and repeating chloroform phase separation. Any residual TRIZol was then removed by washing the interface 2 times with Methanol. The interface was then solubilized in 200 µl of 100mM TEAB; 1mM MgCl2, 1 % SDS and incubated at 95°C for 20 min. The RNA component was digested by addition of 4 µg of RNase A/T1 Mix (Thermo) for 3hrs at 37°C, followed by addition of another 4 µg of RNase A/T1 Mix and incubation at 37°C overnight. The organic Phase was then collected after performing an additional TRIZol / Chloroform phase separation. Acetone precipitation was then performed on the collected organic phases to precipitate the purified proteins.

**Mass spectrometry sample preparation and data acquisition**

For OOPS, acetone precipitated proteins were subjected to in-solution digestion. Briefly, proteins were recovered in 200 µl 2M Urea, 50mM Ammonium Bicarbonate (ABC) and reduced by adding DTT to a final concentration of 10 mM. After 30 minutes of incubation at RT, samples were alkylated by adding 55 mM iodoacetamide and Incubation for another 30 minutes at RT in the dark. Trypsin digestion was then performed using 2 µg of trypsin / sample. The next day, samples were desalted using the Stage Tip procedure (Rappsilber et al., 2003) and recovered in 0.1% TFA, 0.5% Acetic Acid, 2% Acetonitrile (A* buffer) for MS analysis. For total and phospho-proteome analyses, Lysates, in 2-4% SDS, 100mM Tris/HCl pH 7.5, were sonicated with a sonicator bath (Bioruptor Pico - Rm 343) for 10 cycles, and reduced with addition of 100 mM DTT and boiling at 95°C for 10 min. For SILAC samples, Filter Aided Sample preparation (FASP)(Wisniewski et al., 2009) was performed, as described previously (Dermit et al., 2020). For TMT samples, Isobraric Filter Aided Sample Preparation (iFASP) (McDowell et al., 2013) was performed. Briefly, 25 µg (for total proteomics) or 100 µg (for phospho-proteomics) of each total lysate was reduced with 50 mM Bond-Breaker TCEP Solution (Thermo) by boiling at 95°C for 10 min. Reduced samples were then diluted in UA buffer (8 M urea, 100 mM Tris HCl pH 8.8), and transferred to Vivacon 500 Hydrosart filters with a molecular cut-off of 30kDa, before being concentrated by centrifugation at 14,000 g for 20 min. Samples were then washed once with UA buffer through buffer addition to the filter top and concentration, before alkylation with 55 mM iodoacetamide in UA buffer at RT for 30 min in the dark. Samples were then washed three additional times with the UA buffer, before three washes with 100 mM TEAB to reduce the urea concentration. Samples were then trypsin digested overnight at 37°C in a 600 rpm shaking thermomixer, by adding 100 µL of 100mM TEAB supplemented with 25 ng Trypsin per 1 µg of input protein. Next day, TMT 6plex or 10plex label reagents were thawed and dissolved in acetonitrile. Each Sample was then supplemented with 8µg of TMT label per 1µg of input protein, and incubated for 1 hour at 25°C, followed by quenching with 5% hydroxylamine for 30 min at 25°C. Peptides were then eluted by centrifugation at 14,000 g. Two additional elutions were then performed by adding 40 µL of TEAB and centrifugation, plus a final elution with 40 µL of 30% acetonitrile. After combining all individually labeled eluates into one, the pooled mixture was dried with a vacuum concentrator. For SILAC and TMT total proteomics analyses, samples were fractionated into 7 fractions using Pierce™ High pH reverse-phase fractionation kit, according to manufacturer's instructions. Fractions were then dried with vacuum centrifugation before LC-MS/MS analysis. For phospho-proteomics, samples were subjected to TiO phospho-peptide enrichment using GL Sciences TiO enrichment kit, according to manufacturer's instructions. LC-MS/MS analysis was performed on a Q Exactive-plus Orbitrap mass spectrometer coupled with a nanoflow ultimate 3000 RSL nano HPLC platform (Thermo Fisher). Dried peptide mixtures were resuspended in A* buffer. For total proteomics analysis, equivalent of ~ 1 μg of protein was injected into the nanoflow HPLC. For OOPS and phospho-proteomics analysis, ~90% of the total peptide mixture was injected. Samples were resolved at flow rate of 250 nL/min on an Easy-Spray 50cm X 75 μm RSLC C18 column (Thermo Fisher). Each run consisted of a 123 min gradient of 3% to 35 % of Buffer B (0.1% FA in Acetonitrile) against Buffer A (0.1% FA in LC-MS gradient water), and separated samples were infused into the MS by electrospray ionization (ESI). Spray voltage was set at 1.95 kV, and capillary temperature was set to 255°C. MS was operated in data dependent positive mode, with 1 MS scan followed by 15 MS2 scans (top 15 method). Full scan survey spectra (m/z 375-1,500) were acquired with a 70,000 resolution for MS scans and 17,500 for the MS2 scans. For TMT10plex samples, MS2 scans were acquired with 35,000 resolution. A 30 sec dynamic exclusion was applied.

**Mass spectrometry data analysis**

MaxQuant (version 1.6.3.3) was used for all mass spectrometry search and quantifications (Tyanova et al., 2016a). Raw data files were searched against a FASTA file of the Mus musculus proteome, extracted from Uniprot (2016). Enzyme specificity was set to "Trypsin", allowing up to two missed cleavages. False discovery rates (FDR) were calculated using a reverse database search approach, and was set at 1%. Default MaxQuant parameters were used with some adjustments: For TMT experiments, "reporter ion MS2" type option was selected with a reporter mass tolerance of 0.003 Da. For SILAC experiments, "Match between runs" and the "Re-quantify" options were enabled. A minimum ratio count of 2 was also used for protein identifications. All downstream data analyses, such as data filtering, Log transformation, ratio calculation, data normalization, one-sample t-test analysis, category annotation, 1D & 2D annotation enrichment analysis, Fisher's exact test analysis, and data visualizations, were performed in Perseus software (Tyanova et al., 2016b) (version 1.6.2.3). For all annotation enrichments, GO and KEGG annotations were used, with a Benjamini-Hochberg FDR of < 0.02 applied as the cut-off in the adapted Wilcoxon Mann-Whitney test.

**Statistical analysis**

Statistical analyses of the proteomics data were performed using Perseus (version 1.6.2.3) (Tyanova et al., 2016b), as described above. All other statistical analyses were performed using GraphPad PRISM (version 9). Unpaired Student's t-test was applied for western blot images and *in vivo* data analyses. Comparison of all other data was done using Two-way ANOVA.

**Data availability**

All mass spectrometry raw files and their associated MaxQuant output files were deposited on ProteomeXchange Consortium (Vizcaino et al., 2014) via the PRIDE partner repository (http://www.ebi.ac.uk/pride/archive/). In addition, all iCLIP FASTQ files were deposited to the ArrayExpress database (http://www.ebi.ac.uk/arrayexpress).

### Key Resources Table:

| Reagent or resource | Manufacturer | Reference |
|---|---|---|
| **RT-qPCR primers** | | |
| 5’ETS fwd 5’-CTCCCCGTCTTGTGTGTGTCCTCGCCG-3’ | Merck | Custom oligo |
| 5’ETS rev 5’-CCACCCCTTCTCTCACCTCACTCCAGACACCT-3’ | Merck | Custom oligo |
| Actb fwd 5’-CGCCACCAGTTCGCCATGGA-3’ | Merck | Custom oligo |
| Actb rev 5’-TACAGCCCGGGGAGCATCGT-3’ | Merck | Custom oligo |
| **iCLIP 3’ RNA adapter primer** | | |
| (/5Phos/AG ATC GGA AGA GCG GTT CAG AAA AAA AAA AAA/iAzideN/AA AAA AAA AAA A/3Bio/) | Integrated DNA Technologies | Custom oligo |
| **iCLIP Reverse Transcription barcoded primers (5’-3’)** | | |
| /5Phos/ WWW GTGGA NNNN AGATCGGAAGAGCGTCGTGAT /iSp18/ GGATCC /iSp18/ TACTGAACCGC | Integrated DNA Technologies | Custom oligo |
| /5Phos/ WWW TCCGG NNNN AGATCGGAAGAGCGTCGTGAT /iSp18/ GGATCC /iSp18/ TACTGAACCGC | Integrated DNA Technologies | Custom oligo |
| /5Phos/ WWW TGCCT NNNN AGATCGGAAGAGCGTCGTGAT /iSp18/ GGATCC /iSp18/ TACTGAACCGC | Integrated DNA Technologies | Custom oligo |
| /5Phos/ WWW TATTC NNNN AGATCGGAAGAGCGTCGTGAT /iSp18/ GGATCC /iSp18/ TACTGAACCGC | Integrated DNA Technologies | Custom oligo |
| /5Phos/ WWW TTAAA NNNN AGATCGGAAGAGCGTCGTGAT /iSp18/ GGATCC /iSp18/ TACTGAACCGC | Integrated DNA Technologies | Custom oligo |
| /5Phos/ WWW AAATG NNNN AGATCGGAAGAGCGTCGTGAT /iSp18/ GGATCC /iSp18/ TACTGAACCGC | Integrated DNA Technologies | Custom oligo |
| /5Phos/ WWW AAGGT NNNN AGATCGGAAGAGCGTCGTGAT /iSp18/ GGATCC /iSp18/ TACTGAACCGC | Integrated DNA Technologies | Custom oligo |
| /5Phos/ WWW AATAC NNNN AGATCGGAAGAGCGTCGTGAT /iSp18/ GGATCC /iSp18/ TACTGAACCGC | Integrated DNA Technologies | Custom oligo |
| /5Phos/ WWW ACGCA NNNN AGATCGGAAGAGCGTCGTGAT /iSp18/ GGATCC /iSp18/ TACTGAACCGC | Integrated DNA Technologies | Custom oligo |
| /5Phos/ WWW ACTTG NNNN AGATCGGAAGAGCGTCGTGAT /iSp18/ GGATCC /iSp18/ TACTGAACCGC | Integrated DNA Technologies | Custom oligo |
| /5Phos/ WWW AGAGC NNNN AGATCGGAAGAGCGTCGTGAT /iSp18/ GGATCC /iSp18/ TACTGAACCGC | Integrated DNA Technologies | Custom oligo |
| /5Phos/ WWW AGTCT NNNN AGATCGGAAGAGCGTCGTGAT /iSp18/ GGATCC /iSp18/ TACTGAACCGC | Integrated DNA Technologies | Custom oligo |
| **siRNA oligos** | | |
| ON-TARGETplus Non-targeting pool | Dharmacon | D-001810-10-05 |
| ON-TARGETplus Ncl ORF 5’-GCAAAUUCCUAUACAUCUA-3’ | Dharmacon | J-059054-09 |
| ON-TARGETplus Ncl ORF 5’-UGGGAAAAGUAAAGGGAUU-3’ | Dharmacon | J-059054-12 |
| ON-TARGETplus Ncl 3’UTR 5’-GGACAUUCCAAGACAGUAAUU-3’ | Dharmacon | CTM-706612 |
| **Primary antibodies** | | |

| | | |
|---|---|---|
| Anti-GAPDH | Novus Biologicals | NB300-21 |
| Anti-Nucleolin | Abcam | ab22758 |
| Anti-Ras G12D (Mutant Specific) (D8H7) | Cell Signaling | 14429S |
| Anti-p44/42 MAPK (Erk1/2) (137F5) | Cell Signaling | 4695S |
| Anti-Phospho-p44/42 MAPK (Thr202/Tyr204) (p-Erk1/2) (E10) | Cell Signaling | 9106S |
| Anti-Phospho-CK2 Substrate [(pS/pT)DXE] mAb mix | Cell Signaling | 8738S |
| Anti-Fibrillarin (C13C3) | Cell Signaling | 2639 |
| Anti-Phospho-Histone H2A.X (20E3) | Cell Signaling | 9718S |
| Anti-Myc tag (9B11) | Cell Signaling | 2276S |
| Anti-Brdu | Merck | B2531-100UL |
| Anti-NPM1 | Fisher Scientific | 10202223 |
| **Secondary antibodies** | | |
| Rabbit IgG HRP linked | GE Healthcare | NA934 |
| Mouse IgG HRP linked | GE Healthcare | NA931 |
| Cy3-conjugated Donkey Anti-Mouse IgG (H+L) | Jackson ImmunoResearch | 715-165-150 |
| Cy5-conjugated Donkey Anti-Rabbit IgG (H+L) | Jackson ImmunoResearch | 711-175-152 |
| Alexa Fluor 488-conjugated Donkey Anti-Mouse IgG (H+L) | Jackson ImmunoResearch | 715-545-150 |
| Alexa Fluor 647-conjugated Donkey Anti-Rabbit IgG (H+L) | Jackson ImmunoResearch | 711-605-152 |
| DyLight 549-conjugated Donkey Anti-Mouse IgG (H+L) | Jackson ImmunoResearch | 715-505-151 |
| **Chemicals** | | |
| Hoechst 33258 | Thermo Fisher | H3569 |
| NuPAGE LDS Sample Buffer | Thermo Fisher | NP0008 |
| Pierce ECL Plus Western Blotting Substrate | Thermo Fisher | 32132 |
| Luminata Crescendo Western HRP substrate | Fisher Scientific | 10776189 |
| 5-Fluorouridine | Fisher Scientific | 15494529 |
| Crystal violet | Merck | C6158 |
| DTT | VWR | M109 |
| Iodoacetamide | VWR | 786-228 |
| Urea | Merck | U1250—5KG |
| Trypsin | Merck | T6567-1MG |
| TRIzol reagent | Fisher Scientific | 12034977 |
| CX-5461 | Cambridge Biosciences | HY-13323-50MG |
| Silmitasertib | Cambridge Biosciences | 2459-5 |
| Trametinib | Cambridge Biosciences | CAY16292-25 mg |
| SUPERase-In RNase Inhibitor | Thermo Fisher | AM2694 |
| Doxycycline Hyclate | Merck | D9891-10G |
| Chloroform | SLS | 372978-100ML |
| TRIETHYLAMMONIUM BICARBONATE (TEAB) | Fisher Scientific | 15215753 |
| Sodium dodecyl sulfate | Merck | 75746-1kg |
| Magnesium chloride | Severn Biotech Ltd. | 20-xxxx-01 |
| RNAseA, T1 mix | Thermo Fisher | EN0551 |

| | | |
|---|---|---|
| Acetone | Merck | 34850-2.5L |
| Ammonium Bicarbonate | Merck | A6141-500G |
| Trifluoroacetic acid (TFA) | Thermo Fisher | 85183 |
| Acetic Acid | Honeywell | 33209-2.5L |
| Acetonitrile | J.T.Baker | 9012 |
| Tris HCl pH 8.8 | Severn Biotech Ltd. | 20-7900-01 |
| Methanol | Fisher Scientific | M/4056/17 |
| Xylene: 97% | Fisher Scientific | 10784001 |
| IMS (Methylated spirit industrial 74 O.P.) | Fisher Scientific | 11482874 |
| Surgipath Haematoxylin Gill III | Leica Biosystems | 3801542E |
| Surgipath Eosin Y: alcoholic solution | Leica Biosystems | 3801601E |
| 1% Acid Alcohol: Surgipath differentiating solution | Leica Biosystems | 3803650 |
| Formaldehyde solution | Merck | F8775-500ML |
| Nuclease-Free Water | Fisher Scientific | 10526945 |
| Protein G dynabeads | Thermo Fisher | 100.02 |
| RNase I | Thermo Fisher | EN0602 |
| Turbo DNase | Thermo Fisher | AM2238 |
| PNK | NEB | M0201L |
| FastAP thermosensitive alkaline phosphatase | Thermo Fisher | EF0654 |
| RNasin Plus | Promega | N2611 |
| T4 RNA ligase I | NEB | M0204L |
| NEBuffer 2 | NEB | B7002S |
| 5’ deadenylase | NEB | M0331S |
| RecJF endonuclease | NEB | M0264S |
| NuPAGE 4 to 12%, Bis-tris, 1.0mm, mini protein gel, 10-well | Thermo Fisher | NP0321BOX |
| NuPAGE MOPS SDS running buffer | Thermo Fisher | NP0001 |
| Protran nitrocellulose membrane, pore size 0.45μm | Whatman | Z613630 |
| NuPAGE transfer buffer | Thermo Fisher | NP0006 |
| Proteinase K | Thermo Fisher | 25530-049 |
| Phenol:chloroform:isoamyl alcohol | Merck | P3803 |
| 2ml Phase lock gel heavy tube | VWR | 713-2536 |
| Glycoblue | Ambion | 9510 |
| Superscript IV reverse transcriptase kit | Thermo Fisher | 18090010 |
| Exonuclease I | NEB | M0293S |
| Agencourt AMPure XP beads | Beckman Coulter | A63880 |
| CircLigase II ssDNA ligase kit | Epicentre | CL9021K |
| Phusion HF master mix | Thermo Fisher | F531S |
| Novex TBE gel, 6%, 10 well | Thermo Fisher | EC6265BOX |
| SYRB green I 10000X | Thermo Fisher | S7563 |
| Costar SpinX column | Corning Inc. | 8161 |
| 10mm diameter Grade GF/D binder free glass fiber microfiber filter paper circle disc | Whatman | 1823010 |
| Agilent D1000 High sensitivity ScreenTape | Agilent | 5067-5584 |
| Agilent D1000 High sensitivity reagents | Agilent | 5067-5584 |

| | | |
|---|---|---|
| Qubit dsDNA HS assay kit | Thermo Fisher | Q32851 |
| **Tissue Culture reagents** | | |
| iBiDi u-Slide 18 Well flat, ibiTreat, Tissue Culture Treated, Sterile | Thistle Scientific | 81826 |
| Corning Tissue Culture Treated plates and dishes | Corning Inc. | Various products |
| Collagen I (Bovine, pepsinized) | CellSystems GmbH | 5005-100ML |
| Gibco Fetal Bovine Serum | Thermo Fisher | 10437028 |
| Gibco™ DMEM w/High Glucose | Fisher Scientific | 41966-029 |
| Trypsin/EDTA solution | Fisher Scientific | R-001-100 |
| Lipofectamine 2000 Transfection Reagent-1.5 mL | Thermo Fisher | 11668019 |
| Lipofectamine™ RNAiMAX Transfection Reagent | Thermo Fisher | 13778150 |
| Opti-MEM I Reduced Serum Medium-100 mL | Thermo Fisher | 31985062 |
| Gibco DMEM w/High Glucose and w/o Glutamine, Lysine and Arginine (For SILAC) | Fisher Scientific | 12817552 |
| L-Arginine | Merck | A6969-25G |
| L-Lysine | Merck | L8662-25G |
| heavy L-Arginine [U-13C6, U-15N4] | Cambridge Isotopes | CNLM-539-H-0.5 |
| Heavy L-Lysine [U-13C6, U-15N2] | Cambridge Isotopes | CNLM-291-H-0.5 |
| L-Proline | Merck | P0380-100G |
| Gibco Dialyzed Fetal Bovine Serum | Thermo Fisher | 11520646 |
| **Commercial kits and reagent sets** | | |
| TMT6plex™ Isobaric Label Reagent Set | Thermo Fisher | 90061 |
| TMT10plex™ Isobaric Label Reagent Set | Thermo Fisher | 90110 |
| Pierce High pH Reversed-Phase Peptide Fractionation Kit | Life Technologies | 84868 |
| Titansphere TiO phospho-peptide enrichment kit | GL Sciences | 5010-21308 |
| CellTiter-Glo 3D Cell Viability Assay Viability Assay | Promega | G9682 |
| Pierce™ BCA Protein Assay Kit | Thermo Fisher | 23225 |
| Brilliant II SYBR® Green QRT-PCR | Agilent Technologies | 600825 |
| MycoAlert™ PLUS Mycoplasma Detection Kit | Lonza | LT07-705 |
| **Software and algorithms** | | |
| Maxquant | Max Planck Institute | https://www.biochem.mpg.de/5111795/maxquant |
| Perseus | Max Planck Institute | https://www.biochem.mpg.de/5111810/perseus |
| ImageJ | NIH | https://imagej.nih.gov/ij/ |
| Prism | Graphpad | https://www.graphpad.com/scientific-software/prism/ |
| **Other** | | |
| Vivacon 500, 30,000 MWCO Hydrosart | Sartorius | VN01H22 |
| Crl:CD1-Foxn1nu mice | Charles River UK | Strain Code 086 |
| PK20 EMPORE OCTADECYL C18 47MM & | Merck | 66883-U |
| Immobilon-P 26.5 x 3.75m PVDF (0.45µm) | Millipore | IPVH00010 |

### Supplementary figures

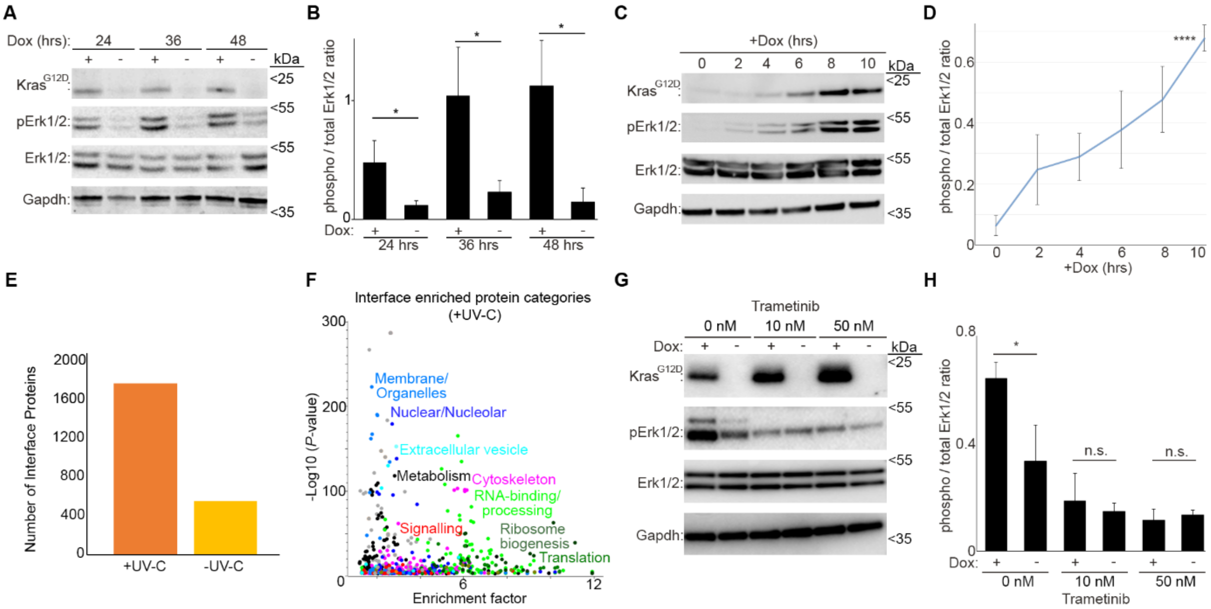

**Figure S1, related to Figure 1: (A)** Doxycycline removal results in loss of Kras$^{G12D}$ expression and ERK activity in iKras PDAC cells. Cells were grown in presence or absence of doxycycline for the indicated amounts of time, before being subjected to lysis and western blotting with the indicated antibodies. **(B)** Quantification of phospho / total Erk1/2 ratio values from (A), as a measure of Erk1/2 kinase activity. A total of 3 independent experiments were quantified (*: $P < 0.05$). **(C)** Addition of doxycycline to doxy-withdrawn cells results in induction of Kras$^{G12D}$ expression and ERK activity in iKras PDAC cells. Cells were grown in the absence of doxycycline for 48 hrs, before its addition for the indicated amounts of time. Cells were then subjected to lysis and western blotting with the indicated antibodies. **(D)** Quantification of phospho / total Erk1/2 ratio values from (C), as a measure of Erk1/2 kinase activity. A total of 3 independent time-course experiments were quantified (****: $P < 0.0001$). **(E)** OOPS-mediated enrichment of proteins in the interface is dependent on UV-C crosslinking. IKras PDAC cells were treated with or without UV-C crosslinking, before lysis in TRIzol and OOPS analysis as in (Queiroz et al., 2019). Interface proteins were then extracted and subjected to mass spectrometry analysis. A total of two biological replicates per condition were analyzed, and the total number of proteins identified in both replicates for each condition were plotted. Enrichment of proteins in the interface was boosted by > 400% upon UV-C crosslinking. **(F)** OOPS specifically enriches RNA-binding proteins in the interface, following UV-C crosslinking. Fisher's exact test analysis of enriched protein categories in the interface of UV-C crosslinked samples from (E) (FDR < 0.02). Each data point represents a category from Gene Ontology (GO) and Kyoto Encyclopedia of Genes & Genome (KEGG) databases, with functionally similar categories highlighted with the same colors. Conventional as well as non-conventional RBPs are significantly enriched in the interface of UV-C crosslinked samples. **(G)** Trametinib inhibits Kras$^{G12D}$ induced ERK activity in iKras PDAC cells. Cells were grown in the absence of doxycycline for 48 hrs, before its addition to the cells for 24 hrs as indicated, with or without the indicated doses of Trametinib. Cells were then subjected to lysis and western blotting with the indicated antibodies. **(H)** Quantification of phospho / total Erk1/2 ratio values from (G), as a measure of Erk1/2 kinase activity. A total of 3 independent experiments were quantified (*: $P < 0.05$; n.s.: non-significant).

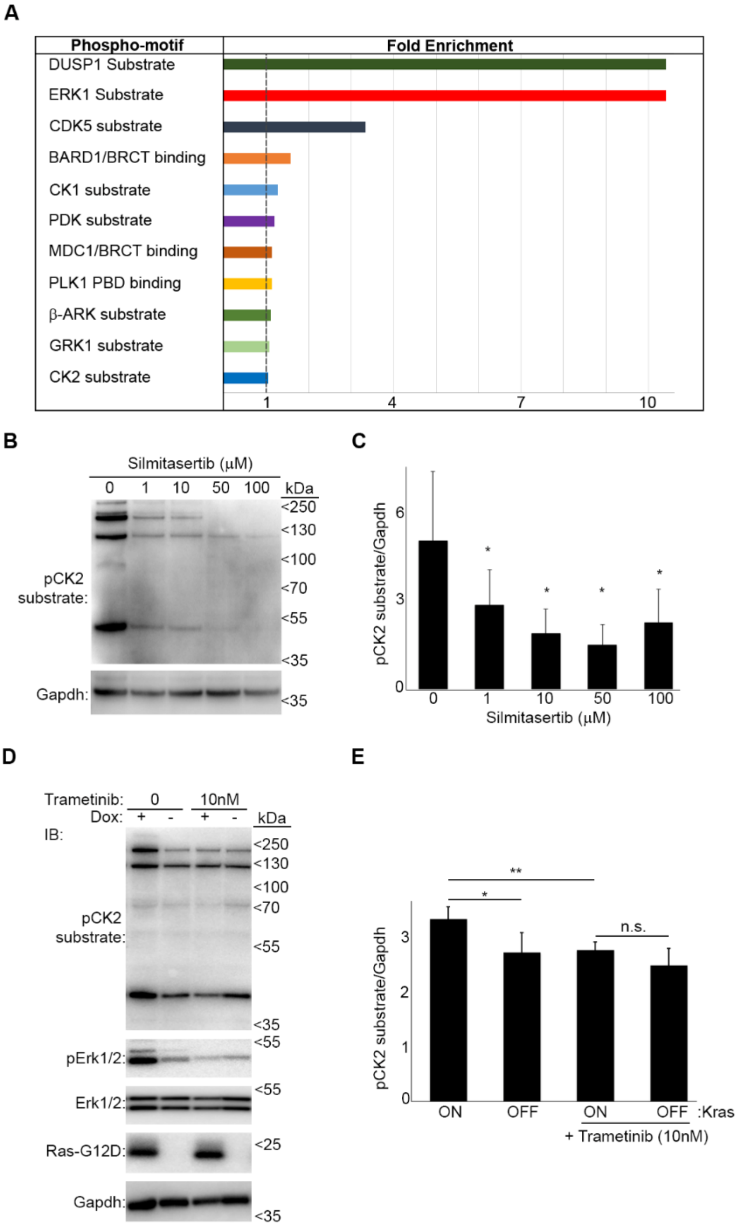

**Figure S2, related to Figure 2: (A)** Phospho-motif analysis of sites which undergo enhanced phosphorylation upon Kras$^{G12D}$ induction in iKras PDAC cells. A total of 11 significantly over-represented motifs amongst the Kras$^{G12D}$ induced phosphorylations (Figure 2A) were identified via Fisher's exact test analysis of Maxquant predicted sequence motifs (FDR <0.02). In addition to the ERK substrate motif, motifs for substrates of CDK5, CK1, CK2, PDK, and GRK/b-ARK kinases were found to be significantly enriched amongst Kras$^{G12D}$ induced phosphorylations. **(B)** A CK2 phospho-substrate antibody mix can be used as an indicator of CK2 activity. IKras PDAC cells grown in presence of doxycycline were treated overnight with the indicated concentrations of Silmitasertib, a specific CK2 inhibitor, before being subjected to lysis and western blotting with the indicated antibodies. **(C)** Quantification of normalized pCK2 substrate levels from (B), as a measure of CK2 kinase activity. A total of 3 independent experiments were quantified (*: $P < 0.05$). **(D)** Kras$^{G12D}$ induction enhances CK2 activity in an Erk1/2-dependent manner. IKras PDAC cells were grown in the absence of doxycycline for 48 hrs, before its addition to the indicated cells, with or without Trametinib (10 nM), for a further 24 hrs. Cells were then lysed and analyzed by western blotting with the indicated antibodies. **(E)** Quantification of normalized pCK2 substrate levels from (D), as a measure of CK2 kinase activity. A total of 3 independent experiments were quantified (**: $P < 0.01$; *: $P < 0.05$; n.s.: non-significant).

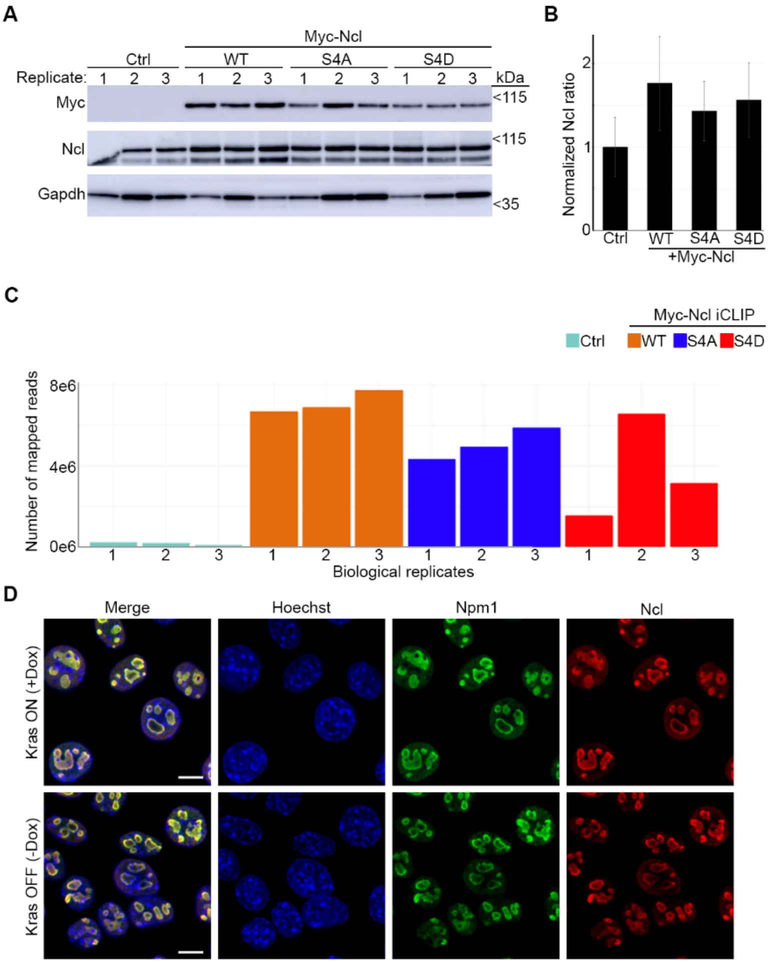

**Figure S3, related to Figure 3: (A)** Assessment of the expression levels of endogenous and ectopic WT, S4A, and S4D Ncl, in iKras PDAC cells that were subjected to iCLIP analysis. IKras PDAC cells were transfected with constructs encoding WT, S4A, and S4D Myc-Ncl, or mock transfected as negative control, before being seeded and grown for 48 hrs. Cells were then UV-C irradiated, lysed, and subjected to iCLIP analysis. Aliquots of the iCLIP input lysates were analyzed by western blotting with the indicated antibodies in parallel. A total of 3 independent replicates per condition were analyzed. **(B)** Quantification of the relative normalized levels of Ncl antibody signal in the Ctrl vs. Myc-Ncl expressing cells from (A). Myc-Ncl transfected cells exhibit total Ncl levels that are around 50% more than those of the Ctrl cells. **(C)** Ncl-bound RNAs are specifically identified in Myc-Ncl iCLIP experiments. Plot of the total number of mapped reads in each replicate of Ctrl vs. Myc-Ncl iCLIP sequencing results. Few reads were identified in the iCLIP sequencing results of Ctrl, as opposed to Myc-Ncl expressing cells. **(D)** Endogenous Ncl is exclusively localized to the Nucleolus of iKras PDAC cells, irrespective of Kras$^{G12D}$ expression. Cells were grown in the absence of doxycycline for 48 hrs, before its addition to the indicated cells for 24 hrs. Doxycycline treated and untreated cells were subsequently fixed and immunostained with an anti-Ncl antibody (red), an anti-Npm1 antibody as a Nucleolar marker (green), and Hoechst (blue) as the Nuclear stain, followed by confocal microscopy analysis. Scale bar = 10 µm.

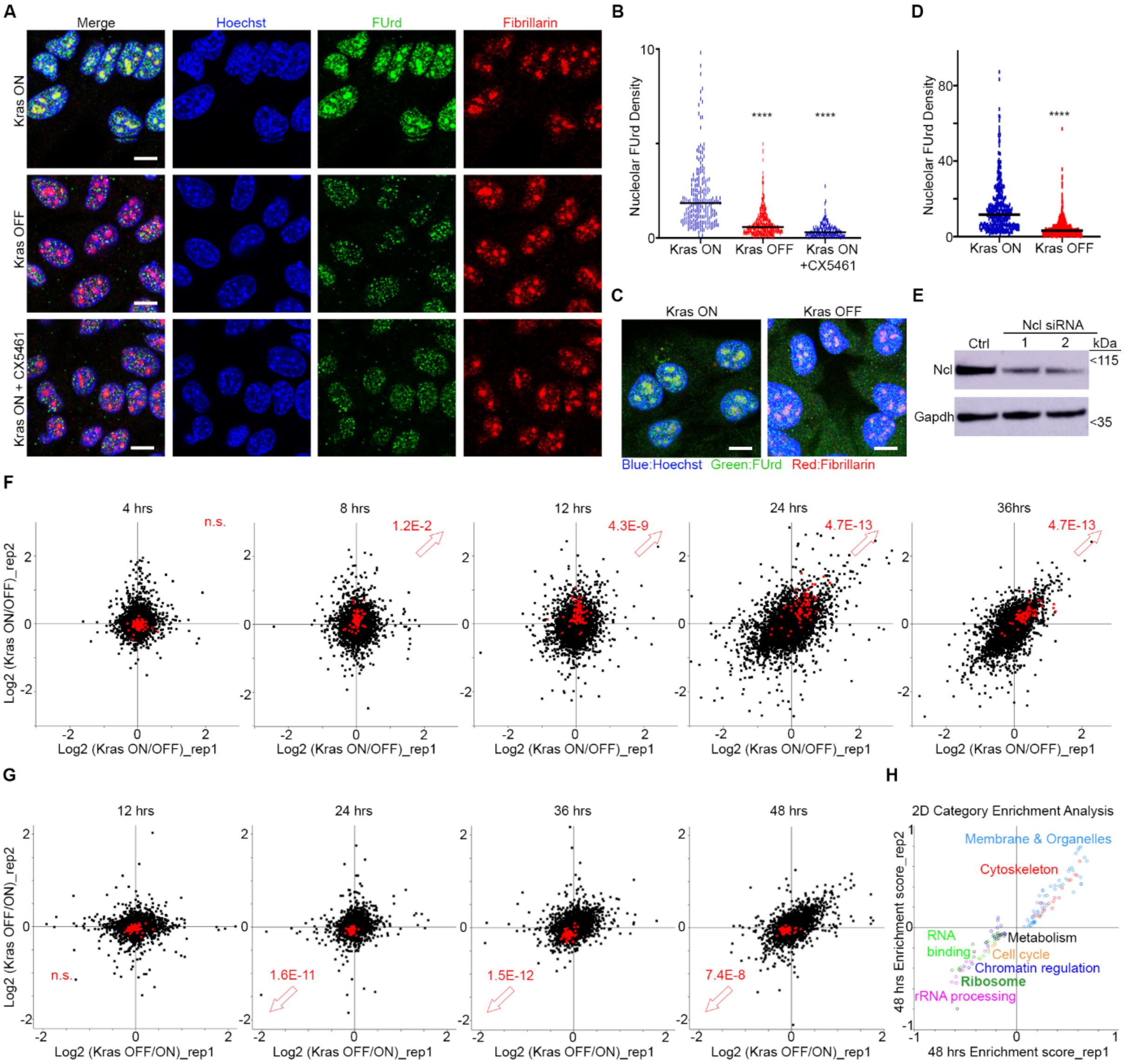

**Figure S4, related to Figure 4: (A)** Induction of Kras$^{G12D}$ expression triggers nascent pre-rRNA synthesis in iKras PDAC cells. Cells were grown for 48hrs in the absence of doxycycline. Kras$^{G12D}$ expression was then induced in the indicated cells (Kras ON) by the addition of doxycycline for a further 24 hrs. The RNA polymerase-I inhibitor (CX-5461) was added to the indicated cells for 30 mins, before all conditions were subjected to pulse labeling with FUrd to visualize RNA synthesis. Cells were then fixed and immunostained with anti-FUrd antibody (green) to visualize nascent RNA, along with anti-Fibrillarin (Fbl) antibody as a Nucleolar marker (red), and Hoechst (blue) as the Nuclear stain, followed by confocal microscopy analysis. Kras$^{G12D}$ induction results in accumulation of nascent RNA in the Nucleoli of iKras PDAC cells, in an RNA polymerase-I dependent manner. Scale bar = 10 µm. **(B)** Quantification of Nucleolar FUrd levels from (A). FUrd fluorescence densities in single nucleoli were quantified from 157-289 individual cells per condition, combined from two independent biological replicate experiments (****: P < 0.0001). **(C)** Removal of Kras$^{G12D}$ expression results in loss of nascent pre-rRNA synthesis in iKras PDAC cells. Cells were grown for 48hrs in the presence or absence of doxycycline, before pulse labeling with FUrd to visualize RNA synthesis. Cells were fixed and immunostained with anti-FUrd antibody (green) to visualize nascent RNA, along with anti-Fibrillarin (Fbl) antibody as a Nucleolar marker (red), and Hoechst (blue) as the Nuclear stain, followed by confocal microscopy analysis. Loss of Kras$^{G12D}$ expression results in abrogation of nascent RNA accumulation in the Nucleoli. Scale bar = 10 µm. **(D)** Quantification of Nucleolar FUrd levels from (C). FUrd fluorescence densities in single nucleoli were quantified from 147-196 individual cells per condition, combined from two independent biological replicate experiments (****: P < 0.0001). **(E)** Validation of siRNA-mediated depletion of Ncl in iKras PDAC cells. Cells were transfected with a non-targeting control siRNA, or two independent siRNAs against Ncl, before being lysed and analyzed by western blotting with the indicated antibodies. **(F)** Induction of $Kras^{G12}$ expression results in accumulation of Ribosomal Proteins (RPs). IKras PDAC cells were grown in the absence of doxycycline for 48 hrs, before its addition to the cells for the indicated amounts of time (Kras ON), or leaving the cells untreated for the same period as control (Kras OFF). Cells were subsequently lysed and subjected to TMT mediated quantitative proteomics (Dataset S9). Log2 of Kras ON/Kras OFF protein ratio values from two replicate experiments were plotted for each time-point, with the ratio values of RPs marked in red. Benjamini-Hochberg corrected P-values of the increase in RPs ratio values is reported on each graph (n.s.: not significant). **(G)** Loss of $Kras^{G12}$ expression results in depletion of RPs. IKras PDAC cells were seeded and grown in the presence (Kras ON) or absence (Kras OFF) of doxycycline for the indicated amounts of time, before lysis and TMT mediated quantitative proteomics (Dataset S10). Log2 of Kras ON/Kras OFF protein ratio values from two replicate experiments were plotted for each time-point, with the ratio values of RPs marked in red. Benjamini-Hochberg corrected P-values of the decrease in RPs ratio values is reported on each graph (n.s.: not significant). **(H)** 2D-annotation enrichment analysis of the 48 hrs time-point data from (G). Each data point represents a functional category from GO and KEGG databases, with similar categories highlighted with the same colors (Dataset S11). After loss of $Kras^{G12}$ for 48 hrs, protein categories related to Ribosome and rRNA processing exhibit significant downregulation, whilst those related to cytoskeleton and membranous organelles show upregulation (FDR < 0.02).

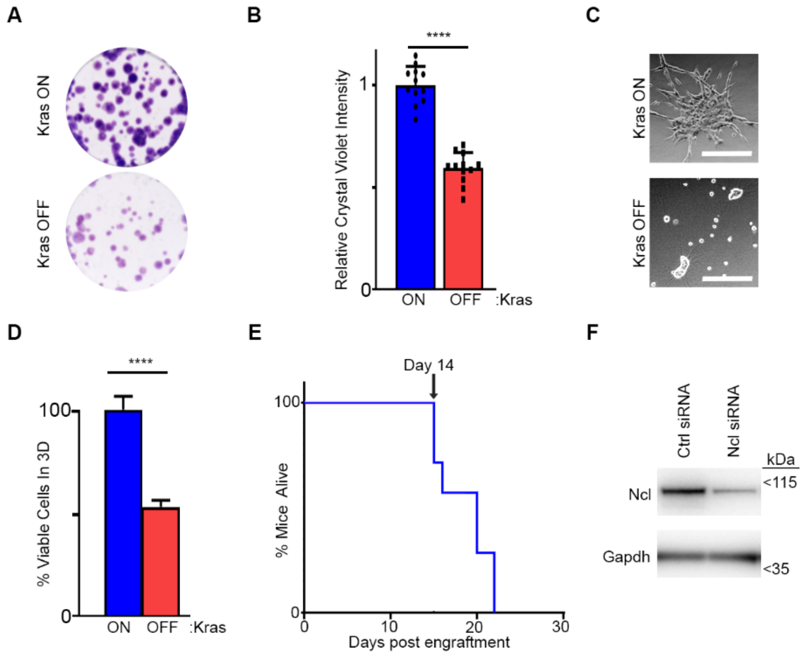

**Figure S5, related to Figure 5: (A)** $Kras^{G12D}$ enhances iKras PDAC cell proliferation in 2D cell-culture. IKras PDAC cells were seeded and subjected to clonogenic assay for 7 days, in the presence (Kras ON) or absence (Kras OFF) of doxycycline. Colonies were visualized by Crystal Violet staining. **(B)** Quantification of Crystal Violet staining levels from (A) (n = 12). (****: P < 0.0001). **(C)** $Kras^{G12D}$ enhances iKras PDAC cell proliferation in 3D cell-culture. IKras PDAC cells were seeded onto 3D Collagen-I gels, with (Kras ON) or without (Kras OFF) doxycycline, and allowed to grow for 48 hrs. Cells were subsequently imaged live by phase contrast microscopy. Scale bar = 200 µm. **(D)** Analysis of the percentage of viable cells in 3D cultures of (C). Cells were subjected to luminescence-based viability assay by CellTiter-Glo to quantify the relative percentage of viable cells (n = 3) (****: P < 0.0001). **(E)** Kaplan-Meier overall survival analysis of nude mice orthotopically engrafted with iKras PDAC cells. (Cohort size = 7). Mice were doxycycline-fed throughout the analysis. Arrow marks the day of the first mortality event. **(F)** Analysis of Ncl expression in iKras PDAC cells that were used for orthotopic xenograft studies in Figure 5E-G.  IKras PDAC cells were transfected with Ctrl and Ncl siRNAs, before injection into the pancreas of nude mice for orthotopic analysis. In parallel, a fraction of the cells from each siRNA treatment was lysed and analyzed by Western blotting with the indicated antibodies.

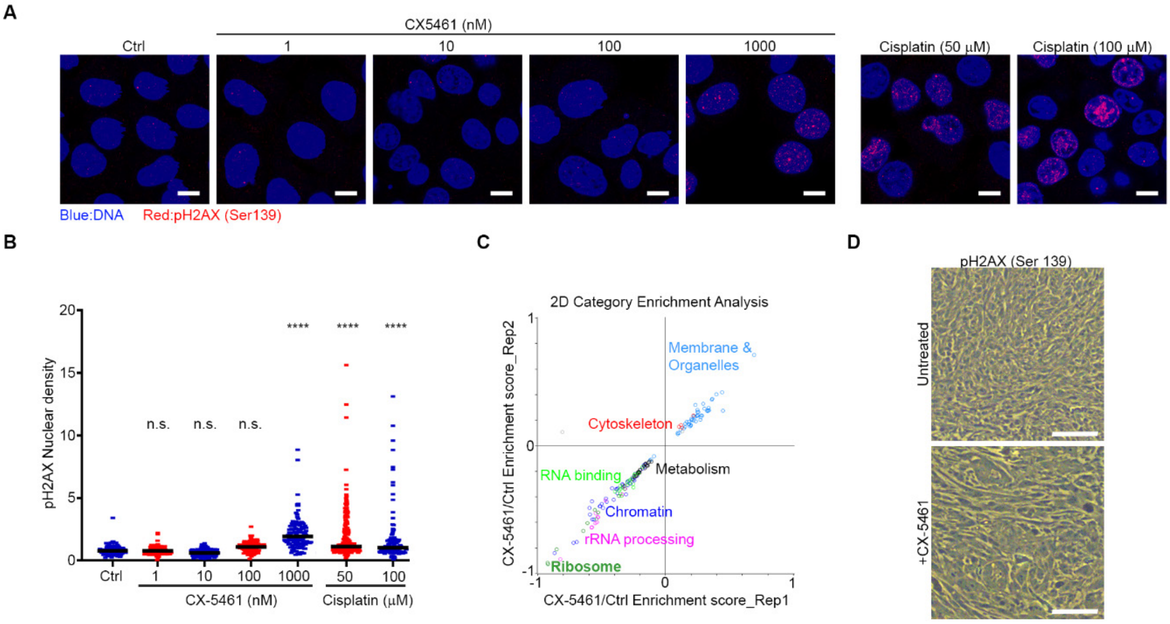

**Figure S6, related to Figure 6: (A)** Dose response analysis of CX-5461 impact on DNA damage. CX-5461 induces DNA damage only at the highest dose tested (1000 nM). IKras PDAC cells grown in the presence of doxycycline were treated overnight with the indicated concentrations of CX-5461. Cells treated with 50 and 100 µM of Cisplatin were used as positive controls. Treated cells were then fixed and immunostained with anti-pH2AX (Ser 139) antibody which marks DNA damage foci (red), and Hoechst as a Nuclear stain (blue), followed by confocal microscopy analysis. Scale bar = 10 µm. **(B)** Quantification of nuclear pH2AX signal intensity from (A). Fluorescence density of pH2AX in single nucleoli were quantified from 55-141 individual cells per condition, combined from two independent biological replicate experiments (****: P < 0.0001; n.s.: not significant). **(C)** Quantification of nuclear pH2AX signal intensity from (A). Fluorescence density of pH2AX in single nucleoli were quantified from 59-141 individual cells per condition, combined from two independent biological replicate experiments (****: P < 0.0001; n.s.: not significant). **(D)** CX-5461 treatment impact on the proteome of iKras PDAC cells mimics $Kras^{G12D}$ removal. IKras PDAC cells grown in presence of doxycycline were treated with or without CX-5461 (100 nM) for 48 hrs, before being lysed and analyzed by TMT-mediated quantitative proteomics. CX-5461 induced changes from two independent replicate experiments were then subjected to 2D-annotation enrichment analysis as in Figure S4H. Each data point represents a functional category from GO and KEGG databases, with similar categories highlighted with the same colors (Dataset S12). Similar to the impact of $Kras^{G12D}$ removal (Figure S4H), protein categories related to Ribosome and rRNA processing exhibit significant downregulation following 48 hrs of CX-5461 treatment, whilst those related to cytoskeleton and membranous organelles show upregulation (FDR < 0.02). No significant change in protein categories related to DNA damage response was detected. **(E)** IHC analysis of tumors from Figure 6I with anti-pH2AX (Ser 139) antibody. No pH2AX signal, indicative of DNA damage, was detectable in tumors from either the control or CX-5461 (50 mg/kg) treated mice. Scale bar = 50 µm.

### Supplementary References

Dermit, M., Dodel, M., Lee, F. C. Y., Azman, M. S., Schwenzer, H., Jones, J. L., Blagden, S. P., Ule, J., and Mardakheh, F. K. (2020). Subcellular mRNA Localization Regulates Ribosome Biogenesis in Migrating Cells. Developmental cell *55*, 298-313 e210.

Dobin, A., Davis, C. A., Schlesinger, F., Drenkow, J., Zaleski, C., Jha, S., Batut, P., Chaisson, M., and Gingeras, T. R. (2013). STAR: ultrafast universal RNA-seq aligner. Bioinformatics *29*, 15-21.

Lee, F. C. Y., Chakrabarti, A. M., Hänel, H., Monzón-Casanova, E., Hallegger, M., Militti, C., Capraro, F., Sadée, C., Toolan-Kerr, P., Wilkins, O.*, et al.* (2021). An improved iCLIP protocol. bioRxiv, 2021.2008.2027.457890.

McDowell, G. S., Gaun, A., and Steen, H. (2013). iFASP: combining isobaric mass tagging with filter-aided sample preparation. J Proteome Res *12*, 3809-3812.

Percipalle, P., and Louvet, E. (2012). In vivo run-on assays to monitor nascent precursor RNA transcripts. Methods in molecular biology *809*, 519-533.

Queiroz, R. M. L., Smith, T., Villanueva, E., Marti-Solano, M., Monti, M., Pizzinga, M., Mirea, D. M., Ramakrishna, M., Harvey, R. F., Dezi, V.*, et al.* (2019). Comprehensive identification of RNA-protein interactions in any organism using orthogonal organic phase separation (OOPS). Nat Biotechnol *37*, 169-178.

Rao, X., Huang, X., Zhou, Z., and Lin, X. (2013). An improvement of the 2^(-delta delta CT) method for quantitative real-time polymerase chain reaction data analysis. Biostat Bioinforma Biomath *3*, 71-85.

Rappsilber, J., Ishihama, Y., and Mann, M. (2003). Stop and go extraction tips for matrix-assisted laser desorption/ionization, nanoelectrospray, and LC/MS sample pretreatment in proteomics. Analytical chemistry *75*, 663-670.

Smith, T., Heger, A., and Sudbery, I. (2017). UMI-tools: modeling sequencing errors in Unique Molecular Identifiers to improve quantification accuracy. Genome Res *27*, 491-499.

Tyanova, S., Temu, T., and Cox, J. (2016a). The MaxQuant computational platform for mass spectrometry-based shotgun proteomics. Nat Protoc *11*, 2301-2319.

Tyanova, S., Temu, T., Sinitcyn, P., Carlson, A., Hein, M. Y., Geiger, T., Mann, M., and Cox, J. (2016b). The Perseus computational platform for comprehensive analysis of (prote)omics data. Nat Methods *13*, 731-740.

Vizcaino, J. A., Deutsch, E. W., Wang, R., Csordas, A., Reisinger, F., Rios, D., Dianes, J. A., Sun, Z., Farrah, T., Bandeira, N.*, et al.* (2014). ProteomeXchange provides globally coordinated proteomics data submission and dissemination. Nat Biotechnol *32*, 223-226.

Wilkins, O. G., Capitanchik, C., Luscombe, N. M., and Ule, J. (2021). Ultraplex: A rapid, flexible, all-in-one fastq demultiplexer. Wellcome Open Res *6*, 141.

Wisniewski, J. R., Zougman, A., Nagaraj, N., and Mann, M. (2009). Universal sample preparation method for proteome analysis. Nat Methods *6*, 359-362.
